# SARS-CoV-2 RdRp uses NDPs as a substrate and is able to incorporate NHC into RNA from diphosphate form molnupiravir

**DOI:** 10.1101/2021.11.15.468737

**Authors:** Maofeng Wang, Cancan Wu, Nan Liu, Fengyu Zhang, Hongjie Dong, Shuai Wang, Min Chen, Xiaoqiong Jiang, Lichuan Gu

## Abstract

The coronavirus disease 2019 (COVID-19) has been ravaging throughout the world for more than two years and has severely impaired both human health and the economy. The causative agent, severe acute respiratory syndrome coronavirus 2 (SARS-CoV-2) employs the viral RNA-dependent RNA polymerase (RdRp) complex for genome replication and transcription, making RdRp an appealing target for antiviral drug development. Here, we reveal that RdRp can recognize and utilize nucleoside diphosphates (NDPs) as a substrate to synthesize RNAs with an efficiency of about two thirds of using nucleoside triphosphates (NTPs) as a substrate. NDPs incorporation is also template-specific and has high fidelity. Moreover, RdRp can incorporate β-d-N4-hydroxycytidine (NHC) into RNA while using diphosphate form molnupiravir (MDP) as a substrate. We also observed that MDP is a better substrate for RdRp than the triphosphate form molnupiravir (MTP).

## Introduction

The pandemic of coronavirus disease 2019 (COVID-19) has severely impacted human health and the global economy since the outbreak in late 2019^1,2^. The causative agent, severe acute respiratory syndrome coronavirus 2 (SARS-CoV-2) is homologous to 2002 SARS-CoV with a genome sequence similarity of 79%^3,4^, forming a sister clade to SARS-CoV and considered a newly β-coronavirus^5^. As of the date of writing, more than 491.75 million infections and more than 6.17 million deaths have been confirmed (https://www.who.int/emergencies/diseases/novel-coronavirus-2019/situation-reports). Although a variety of vaccines have been developed to contain the pandemic, the rapidly spreading mutant strains presses need to develop effective medication or treatment to counter the virus^6–9^.

SARS-CoV-2 is a positive-sense single-stranded RNA virus^5^ with a large genome of approximately 30 kb organized in 14 open reading frames^4,10–12^. Replication of the genome and transcription of genes are all dependent on a protein complex known as RNA-dependent RNA polymerase (RdRp). The RdRp of SARS-CoV-2 contains a catalytic subunit (non-structural protein 12, nsp12) and two accessory subunits (nsp7 and nsp8)^13,14^. Processivity is enhanced by the presence of nsp 7/8 and bound to nsp 12 in a 1:2:1 stoichiometry^15,16^. Due to the significant role of RdRp and the lack of homologues in humans, RdRp has become the most appealing target for anti-coronavirus drug development^17,18^.

To date, the most promising RdRp inhibitors are nucleotide/nucleoside analogs (NA). NA prodrugs are designed to be metabolized to the active 5’-triphosphate form (5’-TP) once inside cells^19^, thus they can compete with endogenous nucleotides for incorporation into newly synthesized viral RNAs, resulting in viral genomes lethal mutation or inactivation^20^. Remdesivir is a broad antiviral drug approved for the treatment of SARS-CoV-2, which can be incorporated into RNA to sterically hinder RdRp, thereby blocking viral RNA synthesis^21^. Other NAs (favipiravir, ribavirin) are integrated into nascent viral RNA at high rates but are not recognized as endogenous nucleotides in subsequent rounds of replication, increasing the mutation rate and resulting in an inviable genome, a process known as “lethal mutagenesis”^22–24^. The recently approved NA antiviral drug molnupiravir significantly increases the frequency of viral RNA mutations and impairs SARS-CoV-2 in both animal models and humans^25,26^

By far none of these RdRp targeting drugs is effective enough to fulfill the need for treatment of COVID-19. To develop a better RdRp targeting drug we need to better understanding of both the properties of RdRp and how these inhibitors function. Previous studies showed that some DNA polymerases^27,28^ and HIV reverse transcriptase^29^ can use deoxynucleoside diphosphates (dNDPs) as a substrate to synthesize DNA, and *Escherichia coli* RNA polymerase^30^ use nucleoside diphosphates (NDPs) to synthesize RNA. Although both dNDPs and NDPs showed very weak activity in these experiments, the data prompted us to test if NDPs are an active substrate for RdRp. In this study, we found that SARS-CoV-2 RdRp can recognize and utilize NDPs as a substrate to synthesize RNAs with an efficiency of about two thirds of using nucleoside triphosphates (NTPs) as a substrate. In addition, we proved that molnupiravir is not only fully functionable in its diphosphate form but also more active than in triphosphate form. This could imply a new strategy for designing COVID-19 treatment nucleoside analogue drugs with more efficacy and fewer side effects.

## Results

### RdRp synthesizes RNA by using both NTPs and NDPs as a substrate

Since RdRp is by far one of the most attractive targets for anti-coronavirus drug development, a thorough characterization of RdRp polymerase activity will, without doubt, greatly contribute to discovering the effective strategy for drug screening^17,18^. The RdRp of SARS-CoV-2 was purified by assembling nsp12, nsp7 and nsp8 (Fig.1a). Based on previous research, SARS-CoV-2 RdRp polymerase activity was measured by conducting a conventional RNA elongation assay^31^. When NTPs was added, elongation catalyzed by RdRp complex occurred on the RNA duplex, resulting in an intact double-stranded RNA product (Fig.1c).

**Fig.1.**
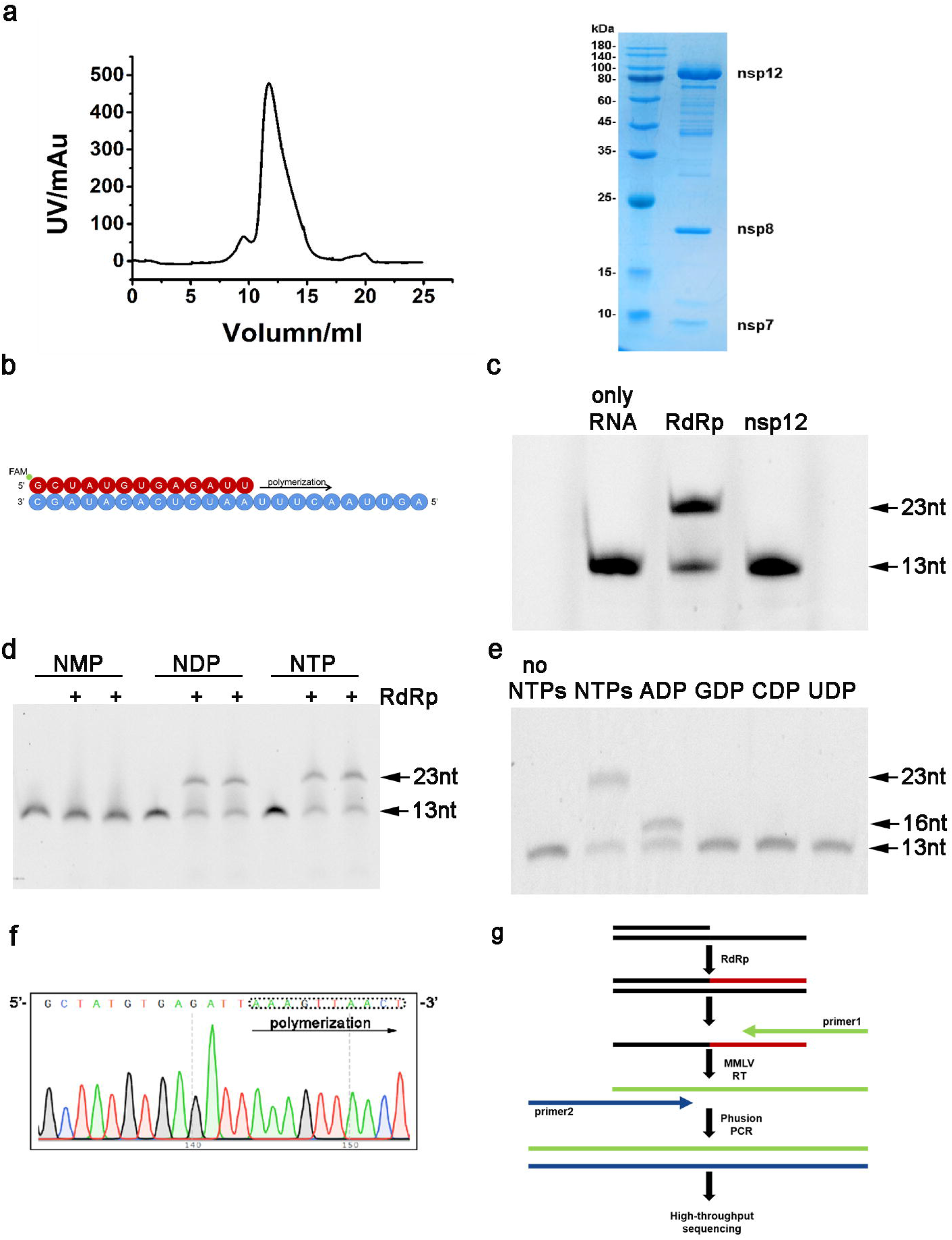
RdRp uses both NTPs and NDPs as a substrate. **a** Size-exclusion chromatogram of the SARS-CoV-2 nsp12-nsp7-nsp8(RdRp) complex. RdRp was also characterized by SDS-PAGE. **b** The 13+23 nt RNA template-product duplex. The direction of RNA extension is shown. The color of the depicted circles indicates the experimental design: blue, RNA template strand; red, RNA product strand. The 5’end of the RNA product contains a FAM fluorescent label. **c** Incubation of the RdRp with RNA duplex and NTPs leads to RNA extension. Nsp12 alone has almost no polymerase activity, and requires nsp7 and nsp8 to form an RdRp complex to have polymerase activity. **d** RdRp was incubated with the 13+23 nt duplex in the presence of NMPs, NDPs or NTPs respectively. These are showing duplicate experiments performed with different batches RdRp. The positions of the template RNA and the full-length extension product are indicated on denaturing gel. **e** Electrophoretic separation of reaction products of single nucleoside diphosphates as substrates. The positions of the original RNA and extension products are indicated. **f** Part of the Sanger sequencing chromatogram of the RT-PCR products corresponding to the extended 10 nt RNA on the 13+23 nt RNA duplex with NDPs as substrate. The dash line circle represents the 10 nt extension. **g** Schematic of high-throughput sequencing of RdRp reaction products.

To test if RdRp uses NDPs or NMPs as a substrate, we performed an RNA extension assay by using NTPs, NDPs or NMPs as substrate, respectively. The result revealed that RdRp synthesized RNA products of the same length regardless of NTPs or NDPs as substrate (Fig.1d). To examine the template specificity of this RNA synthesis, we challenged 13+23 nt RNA with various NDPs (Fig.1b). Only the cognate ADP promoted RNA extension, indicating that NDPs incorporation is template-specific (Fig.1e).

This result also raised the question of whether the RNA made from NDPs has the right sequence. Hence, two experiments were performed: single-clonal sequencing of ligated plasmids, and high-throughput sequencing of PCR products. First, the cDNA was synthesized through reverse transcription of the RNA products, and then amplified by 10 cycles of PCR with primers containing the restriction sites of XhoI and BamHI. PCR products were then ligated into the vector pET-15b and used to transform *E. coli* DH5α strain. Plasmids extracted from the positive clones were sequenced by Sanger sequencing. The sequencing results indicated that the RNA synthesized from NDPs has the right sequence (Fig.1f). Second, the cDNA amplified by 10 cycles of PCR was sequenced by high-throughput sequencing (Fig.1g). At least 50,000 sequences were obtained for each product (SI). The ADP, GDP, CDP and UDP (purchased from Sigma) used in these experiments were characterized by mass spectrometry respectively to make sure these NDPs were not contaminated by NTPs (S1).

### NDPs are nearly two thirds as efficient as NTPs as a substrate of RdRp

Previous studies indicated that although some DNA polymerases and RNA polymerases are able to use diphosphate form substrate. The activity of the diphosphate form substrate, however, is much lower than that of the triphosphate form substrate^28–30^. To determine if this is also the case for RdRp we performed a kinetics study by using NTPs and NDPs as a substrate respectively. The 13+23 nt RNA duplex was used as a template and the elongation of product over time was observed (Fig.2a). Kinetics curves indicated that the relative incorporation efficiency of NDPs is about two thirds (67.94% ± 2.66%) of that of NTPs (Fig.2b). These data suggested that the incorporation rate for NDPs is slower than that of NTPs. Subsequently, we decided to test if all four NDPs equally contribute to the slower incorporation rate. A 13+26 nt RNA duplex was synthesized for the assay testing the incorporation efficiency for each NDP (Fig.2c). Each reaction system contains three NTPs and the NDP of interest (Fig.2d). Unexpectedly, our data showed that substitution of any single NTP by NDP has no observable effect on the efficiency of RNA synthesis (Fig.2e). This may imply that the speed of RNA synthesis is only affected by continuous NDP incorporation but not the intermittent insertion.

**Fig.2.**
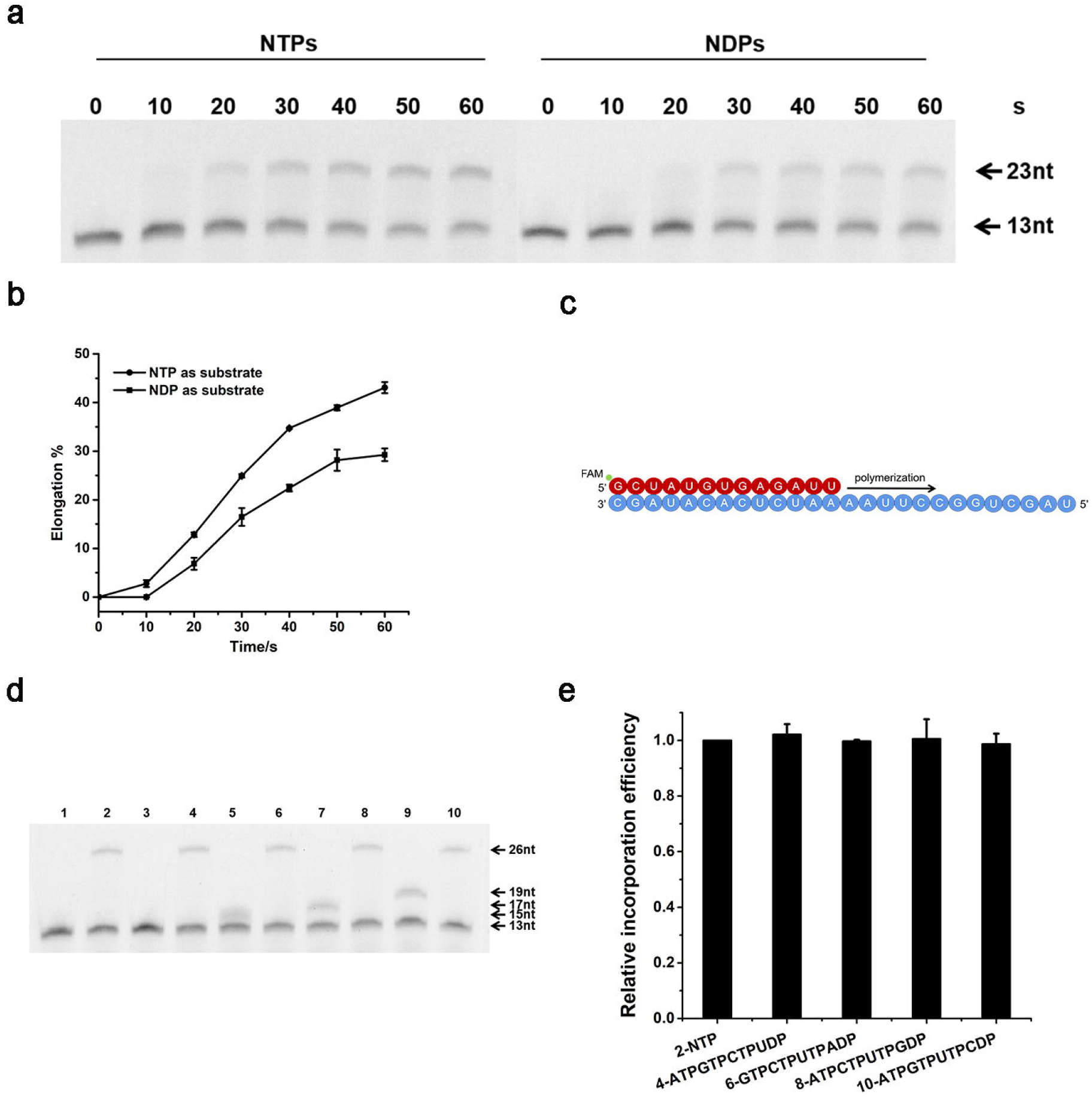
The activities of NDPs and NTPs are comparable. **a** Primer-extension reactions on 13+23 nt RNA templates sampled at indicated time points. **b** Quantification of elongation products for (**a**). The experiment was performed in triplet. **c** The 13+26 nt RNA template-product duplex. The direction of RNA extension is shown. The 5’end of the RNA product was labeled by FAM. **d** RdRp catalyzed reactions containing different substrates. Lane 1: no substrate; lane 2: NTPs; lane 3: ATPGTPCTP; lane4: ATPGTPCTP UDP; lane5: GTPCTPUTP; lane6: GTPCTPUTP ADP; lane7: ATPCTPUTP; lane8: ATPCTPUTP GDP; lane9: ATPGTPUTP; lane10: ATPGTPUTP CDP. The positions of the 13+26 nt RNA and extension products are indicated. **e** Quantification of elongation products for (**d**).

### SARS-CoV-2 RdRp incorporates NHC into RNA by using diphosphate form molnupiravir as a substrate

Since RdRp has become one of the most important targets for the development of antiviral drugs, many nucleoside analogue drugs have been constructed to target RdRp for COVID-19 treatment^24,32–34^. Among all these chemicals reported by far, molnupiravir has been regarded as a promising drug candidate. The molecular mechanism of molnupiravir was determined recently. Once ingested by patients, molnupiravir undergoes stepwise phosphorylation to yield the active nucleoside triphosphate analogue (β-d-N4-hydroxycytidine (NHC) triphosphate, or MTP). The RdRp then uses MTP as substrate and incorporates NHC monophosphate into RNA at the position where the nucleotide in the template is G or A, resulting in “error catastrophe” and virus death^26,35,36^ (Fig.3a).

**Fig.3.**
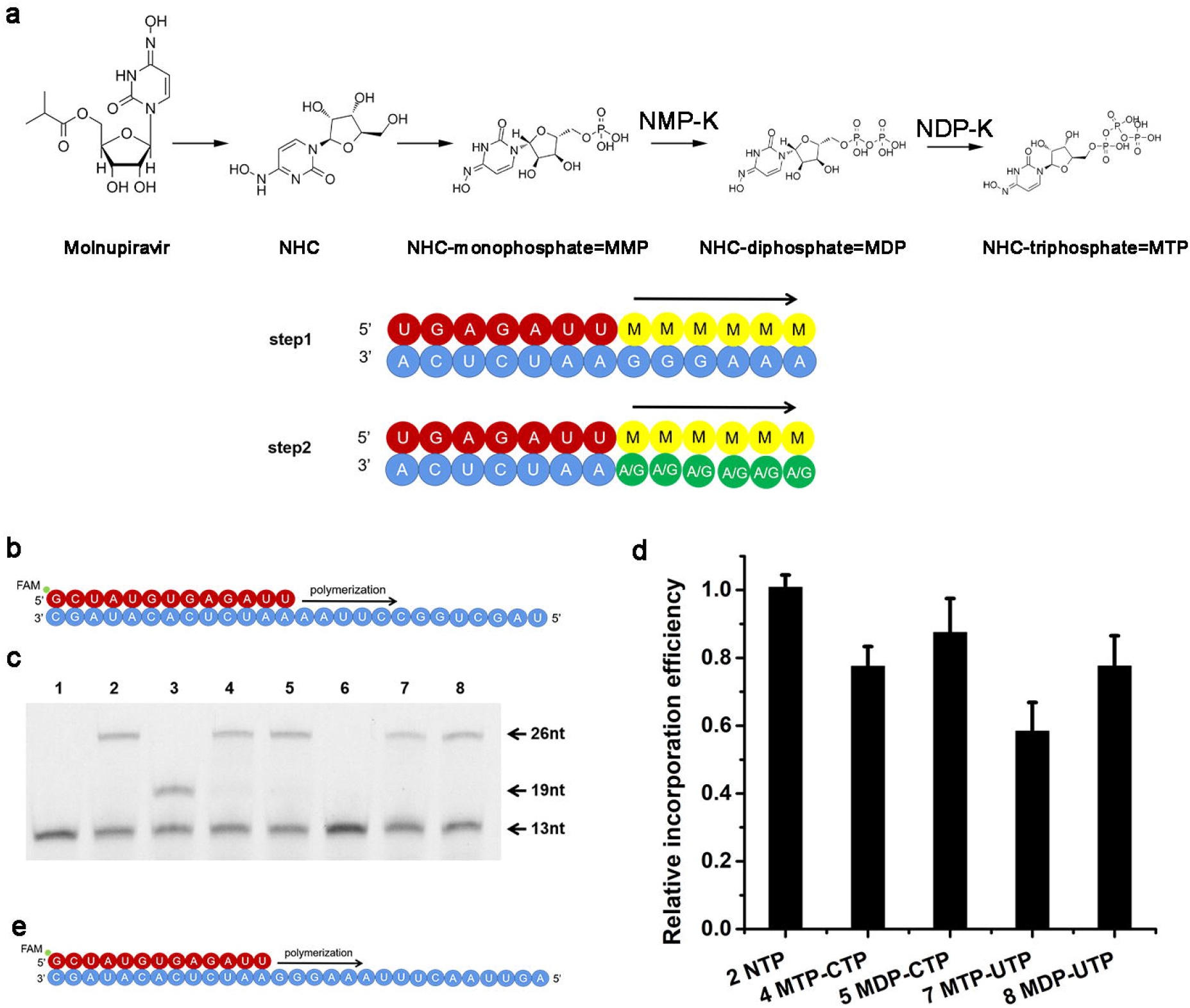
RdRp incorporates NHC into RNA with MDP as a substrate. **a** Metabolism of nucleoside analogues molnupiravir in host cell. A two-step model of molnupiravir-induced RNA mutagenesis. **b** The 13+26 nt RNA allows for RNA extension by 13 nucleotides. NHC monophosphate can be incorporated into growing RNA instead of C or U. **c** RdRp catalyzed reactions containing different substrates. Lane 1: no substrate; lane 2: NTPs; lane 3: ATPGTPUTP; lane4: ATPGTPUTP MTP; lane5: ATPGTPUTP MDP; lane6: ATPGTPCTP; lane7: ATPGTPCTP MTP; lane8: ATPGTPCTP MDP. The positions of the 13+26 nt RNA and extension products are indicated. RNA elongation stalls at the expected positions when the cognate NTP is absent from the reaction. **d** Quantification of elongation products for (**c**). **e** The 13+29 nt RNA allows for RNA extension by 16 nucleotides. NHC monophosphate can pair with G or A for incorporation into growing RNA.

Although the mechanism of molnupiravir has been well established, our finding of-RdRp’s ability to utilize NDPs as a substrate still raises a question: is NHC diphosphate (MDP) also an active form and recognized by RdRp as a substrate? To address this question, the 13+26 nt RNA duplex was used to perform the RNA extension assay with NDPs and MDP as the substrates in comparison with the extension using NTPs and MTP as the substrates. The 13+26 nt RNA duplex allows ten nucleotides (nt) of extension. The 13+26 nt RNA duplex contanis two adjacent G and two adjacent A, allowing NHC incorporation into the RNA (Fig.3b). In this case, when cytidine triphosphate (CTP) or uridine triphosphate (UTP) is replaced by MDP, the incorporation of M does not hinder the incorporation of the next subsequent nucleotide, and RdRp still completes the extension reaction. Similar results were obtained using MTP as the substrate (Fig.3c). Therefore, it is reasonable to speculate that both MTP and MDP are active forms of monulpiravir in humans.

### MDP is a better substrate than MTP

Since both MTP and MDP are active substrates of RdRp we performed an assay to figure out which one is the better substrate. By observing their efficiency in reaction, we found that when the template is G, the relative incorporation efficiency of MTP is 0.78, while MDP is 0.87 in comparison to CTP. When the template is A, the relative incorporation efficiency of MTP is 0.58, while MDP is 0.77 in comparison to UTP (Fig.3d). In order to obtain a reliable result, the experiment was repeated six times with two batches of RdRp. The results indicated MDP is the better substrate for RdRp in comparison to MTP. MTP and MDP were characterized by HPLC and mass spectrometry to make sure they are the right pure compounds (S2).

Next, we tried to determine if MDP is also more efficient in introducing mutations than MTP. The products of RdRp-catalyzed extension reactions containing MTP or MDP using 13+29 nt RNA as a template (Fig. 3e) were reverse-transcribed and PCR-amplified, and the number of mutations was counted by high-throughput sequencing. The statistical results were shown in Table 1. Similar mutation ratios were found when the same concentrations of MTP and MDP were added. Higher ratio of MTP or MDP to CTP associates with higher mutation rate, with MDP resulting in slightly more mutations.

**Table.1.**

| substrate | NTP | 50 μM CTP<br>50 μM MTP | 50 μM CTP<br>50 μM MDP | 5 μM CTP<br>50 μM MTP | 5 μM CTP<br>50 μM MDP | 0.8 μM CTP<br>50 μM MTP | 0.8 μM CTP<br>50 μM MDP |
| --- | --- | --- | --- | --- | --- | --- | --- |
| all sequences | 67883 | 64316 | 61673 | 59588 | 70906 | 59129 | 56214 |
| no mutation<br>(TCCCTTT) | 67105 | 63626 | 61054 | 58386 | 69443 | 57831 | 54945 |
| % of no mutation | 98.85% | 98.93% | 99.00% | 97.98% | 97.94% | 97.80% | 97.74% |
| mutations* | 778 | 690 | 619 | 1202 | 1463 | 1298 | 1269 |
| % of mutation | 1.15% | 1.07% | 1.00% | 2.02% | 2.06% | 2.20% | 2.26% |

\*The mutation resulting from the incorporation of M is shown as: TCCCTTC, TTCCTTT, TACCTTT, TCCCCTT, TCACTTT, TCCCTCT, TCCCTAT etc.

### NDPs are a good substrate for SARS-CoV-2 RdRp but not other virus RNA polymerases

To test if other virus RNA polymerases also use NDPs as a substrate, we purified SARS-CoV RdRp, NS5 protein of ZIKA virus and T7 RNA polymerase (RNAP) and measured their activities when NDPs were used as s substrate. Results show that amount these RNA polymerases tested SARS-CoV-2 RdRp had the highest activity when using NDPs as a substrate. SARS-CoV RdRp could also use NDPs as a substrate, but the product band seam a double RNA strand shorter than expectation implying stalled extension. NS5 of ZIKA virus did not produce a well-defined product band when using NDPs as a substrate (Fig.4a). The transcription of T7 RNAP was measured based on fluorescence technology. We found that except for the case in which ATP was replaced by ADP and weak transcription occurred, any other NTP replaced by NDP would inhibit the transcription completely. (Fig.4b).

**Fig.4.**
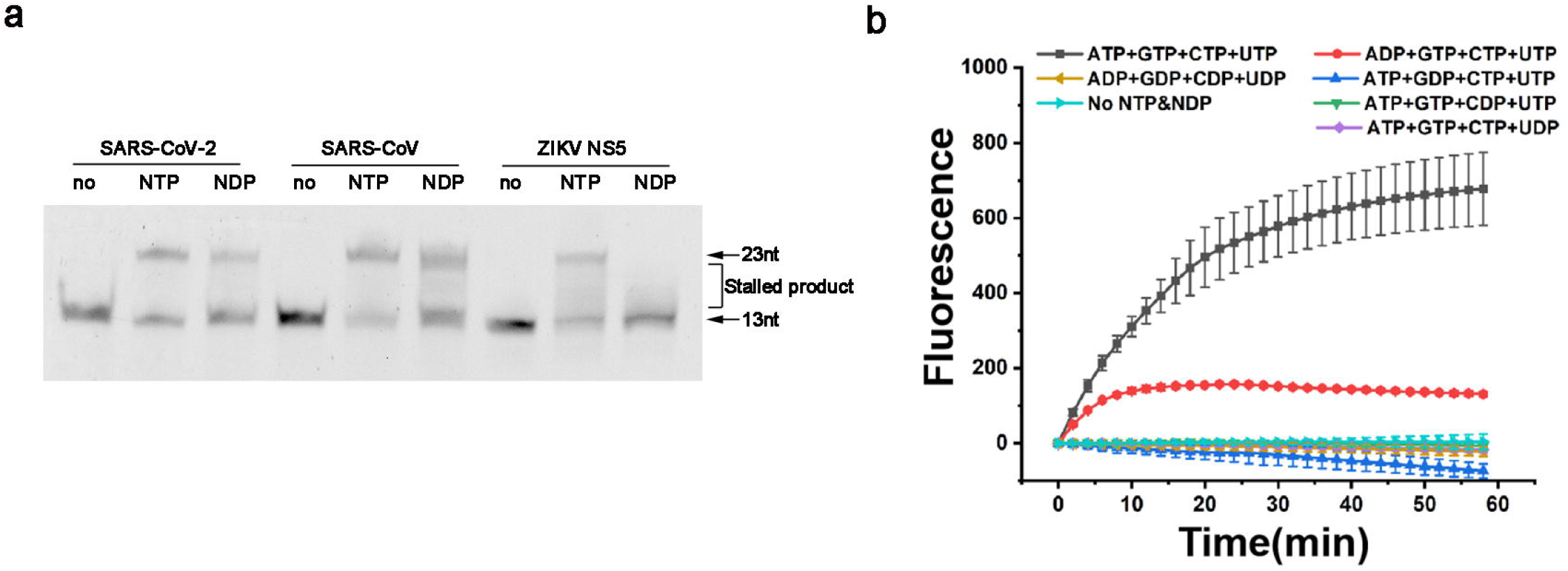
Substrate properties of other RNA viruses **a** RdRp of SARS-CoV-2 and SARS-CoV and NS5 of ZIKV were incubated with the 13+23 nt RNA duplex in the presence of NTPs or NDPs respectively. The positions of the template RNA and the full-length extension product are indicated. **b** The transcription of T7 RNA polymerase was measured by monitoring the signal of a fluorescent probe. Fluorescent signal continuously changed when transcription occurred, indicating the presence of RNA product. However, when NDPs were used as a substrate, the fluorescent signal remained unchanged, indicating the lack of RNA product. Substitution of ATP by ADP led to weak transcription, any other NTP replaced by NDP would inhibit the transcription completely.

## Discussion

Since its outbreak in late 2019, the COVID-19 pandemic has been plaguing people for two years with no sign of abating^3,11^. Although many kinds of vaccines have been developed to suppress the pandemic, they are not sufficient to withhold the spread of SARS-CoV-2^37^. The development of potent antivirals against SARS-CoV-2 is still an urgent need^6,38^.

Since RNA-dependent RNA polymerase (RdRp) is critical for viral genome replication and transcription and conserved in RNA virus species, RdRp has become one of the most appealing targets for antivirals development^17,31,38,39^. Thoroughly characterization of RdRp both structurally and biochemically would no doubt be beneficial to drug development.

The substrate usage of RdRp is astonishing. To the best of our knowledge, most RNA polymerases, including DNA-dependent and RNA-dependent RNA polymerases, use nucleoside triphosphates (NTPs) as a substrate to synthesize RNA under the guidance of a template strand. We found that SARS-CoV-2 RdRp synthesizes RNA with comparable efficiency using NTPs and NDPs as a substrate, and other RNA viruses that can utilize NDPs as a substrate have also been found. We speculate that the ability to use NDPs as a substrate may be an advantage of RNA virus during infection. During later stages of infection, host cells may not be able to produce enough ATP and other NTPs. By using NDPs as a substrate, SARS-CoV-2 can continue the process of genome replication and assembly of progeny viruses, while most metabolic activities stop.

It has long been a general knowledge that to become active against virus nucleoside analogue drugs must undergo stepwise addition of phosphate groups to become the triphosphate form. In order to produce this outcome, this type of antivirals must be recognized by three kinds of kinases^40^. This raises a concern that the triphosphate form of antivirals may be incorporated into host mRNA^36^. Even worse, mutagenic ribonucleoside analogs could be reduced into the 2’-deoxyribonucleotide form by host ribonucleotide reductase and then incorporated into DNA^35,36^. Our finding that RdRp also uses MDP as a substrate could largely resolve these concerns. Since the diphosphate form is also active, there would be no issue regarding whether the prodrugs can become triphosphate form inside host cells. Furthermore, nucleoside diphosphate analogues, which can evade nucleoside diphosphate kinase, would cease to have the possibility of being incorporated into mRNA or DNA. The best strategy is to use the membrane-permeable nucleotide diphosphate analog for COVID-19 treatment^40,41^. The advantage of this strategy is that the drug is not only able to bypass all the phosphorylation steps and has no risk of incorporation into host mRNA or DNA (Fig.5).

**Fig.5.**
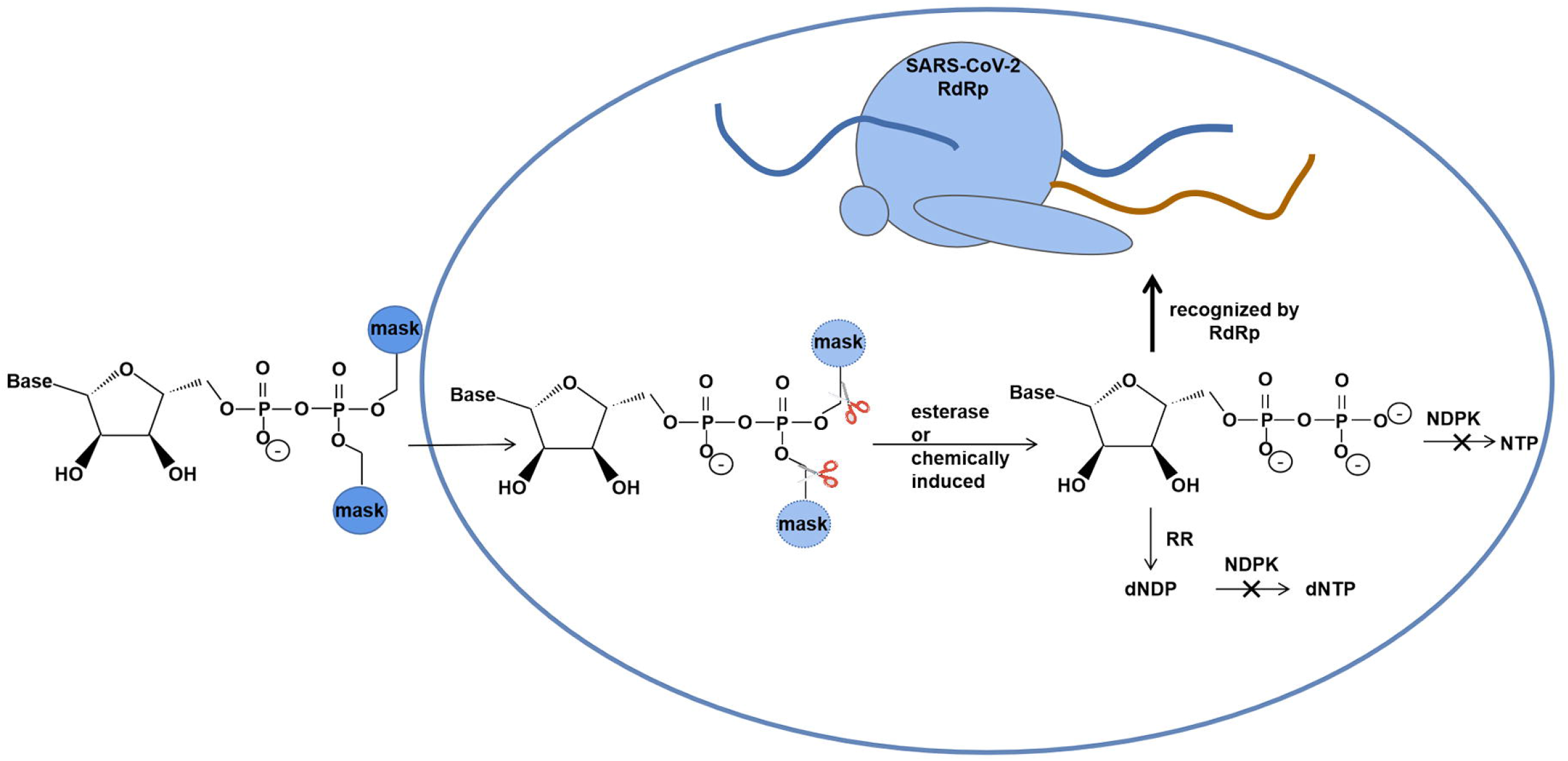
Nucleotide diphosphate analog prodrugs designed by our strategy would have less potential to cause side effects. The terminal phosphate group (β-phosphate) is modified by lipophilic masks to give the prodrug more membrane permeability. Inside cell the masks are removed by enzymatic or chemical reactions. The ribose moiety and the base group should be designed in such a way that the diphosphate form drug can evade NDPK thus eliminating the risk of being incorporated into host mRNA or DNA (Note: NDPs and dNDPs are phosphorylated by the same NDPK).

## Materials and Methods

### Protein expression and purification

Preparation of SARS-CoV-2 RdRp, composed of nsp12, nsp7 and two copied of the nsp8 subunits, was carried out as described in ref.^31^. The SARS-CoV-2 *nsp12* and *nsp7* gene were cloned into a modified pET-21b vector with the C-terminus possessing a 6×His-tag. The *nsp8* gene was cloned into the modified pET-32a vector with the N-terminus possessing a trx-His6-tag and PreScission Protease site. The *nsp12*-pET-21b and *nsp8*-pET-32a plasmids were transformed into *E. coli* BL21 (DE3) and the transformed cells were cultured at 37 °C in LB with a final concentration of 100 μg/ml ampicillin. The *nsp7*-pET-21b plasmid was transformed into *E. coli* Rosetta-gami2 (DE3) and the transformed cells were cultured at 37 °C in LB contanning a final concentration of 100 μg/ml ampicillin and 25 μg/ml chloramphenicol.

Bacterial cultures were incubated by shaking at 200 rpm, 37 °C to an OD600 of 0.8, then the temperature was lowered to 16 °C and a final concentration of 0.2 mM of isopropyl β-D-1-thiogalactopyranoside (IPTG) was added to induce protein expression for 20 h. Subsequently, the cells were harvested by centrifugation at 5000 ×*g* for 15 min at 4 °C. The pellet was resuspended in lysis buffer containing 25 mM TrisHCl pH 8.0, 250 mM NaCl, 4 mM MgCl_2_, 10% Glycerol and homogenized with a high-pressure cell disrupter at 4 °C. The lysate was centrifuged at 25,200 ×*g* for 50 min at 4 °C, and the supernatant was then loaded onto Ni-NTA affinity chromatography column for purification. Nsp12, nsp7 were eluted with elution buffer (25 mM TrisHCl pH 8.0, 200 mM NaCl, 4 mM MgCl_2_, 250 mM Imidazole). Nsp8 was subject to on-column tag cleavage by PPase (PreScission Protease) and then eluted by the lysis buffer. Next, all proteins were purified by ion exchange chromatography (Source 15Q HR 16/10, GE Healthcare) and size exclusion chromatography (Superdex 200 10/300GL, GE Healthcare) with 25 mM TrisHCl pH 8.0, 200 mM NaCl, 4 mM MgCl_2_.

For nsp12-nsp7-nsp8 complex assembly, protein samples were mixed with the molar ratio of nsp7:nsp8:nsp12=2:2:1 at 4 °C overnight. The incubated RdRp complex was then concentrated with a 100 kDa molecular weight cut-off centrifugal filter unit (Millipore Corporation) and then further purified by size exclusion chromatography using a Superdex 200 10/300 GL column in 25 mM TrisHCl pH 8.0, 200 mM NaCl, 4 mM MgCl_2_. SARS-CoV RdRp was purified by the same procedure. The NS5 RdRp domain of ZIKV was purified as previously described^42^.

### Preparation of fluorescently labeled RNA for polymerase activity

RNA template-product duplex was designed according to the published SARS-CoV-2 RNA extension assays^31^. FAM labeled 13 nt oligonucleotide with the sequence of FAM - GCUAUGUGAGAUU and the 23 nt complementary RNA strand with the sequence of AGUUAACUUUAAUCUCACAUAGC were synthesized for polymerase activity assay. A 26 nt RNA strand with the sequence UAGCUGGCCUUAAAAUCUCACAUAGC was synthesized for substrate efficiency assay. And a 29 nt RNA strand with the sequence AGUUAACUUUAAAGGGAAUCUCACAUAGC were used for the detection of mutation rates due to the incorporation of MTP and MDP into RNA. All unmodified and 5’ FAM-labeled RNA oligonucleotides were purchased from Tsingke Biotechnology Co.,Ltd. The RNA strands were mixed in equal molar ratio in DEPC water, annealed by heating it to 95 °C for 10 min and gradually cooling to room temperature to make the RNA duplexes.

### RdRp polymerase activity assays

DEPC-treated water was used in the preparation of all solutions. RdRp at final concentration of 2 μM was incubated with 200 nM 13+23 nt RNA duplex and 50 μM NTPs (or NDPs / NMPs) in a 20 μl reaction buffer containing 20 mM TrisHCl pH8.0, 10 mM KCl, 10 mM MgCl_2_, 0.01% Triton-X100, 1 mM DTT for 1 min at 37 C^31^, the reactions were stopped with 2 × stop buffer (10 M urea, 50 mM EDTA). Eventually, glycerol was added to a concentration of 6.5%. 10 μl RNA product for each reaction was resolved on 20% denaturing polyacrylamide-urea gels and imaged with a Tanon-5200 Multi Fluorescence Imager. The experiments described were performed in triplet, unless specified otherwise.

### RNA extension assays with NDPs as substrate

RdRp of different viruses at final concentration of 2 μM was incubated with 200 nM 13+23 nt RNA duplex and 50 μM NTPs (or NDPs/NMPs/ADP/GDP/CDP/UDP) in a 20 μl reaction buffer containing 20 mM TrisHCl pH8.0, 10 mM KCl, 10 mM MgCl_2_, 0.01% Triton-X100, 1 mM DTT for 1 min at 37 °C.

### Sanger sequencing

The RNA product, primer1 (CCGCTCGAGCGGAGTTAACTTT) and dNTPs were gently mixed at 65 °C for 10 min. The mixture was cooled down to 24 °C then 10× MMulV buffer and MMulV reverse transcriptase were added and incubated for 2 h to obtain cDNA. Thereafter, the reverse transcriptase was inactivated at 70 °C for 15 min. Primer2 (CGCGGATCCGCGGCTATGTGAGAT) was then added, and the product was amplified by PCR catalyzed by the high-fidelity DNA polymerase pfu-Phusion. The PCR product was digested with XhoI and BamHI and ligated into pET-15b vector, and transformed into *E-coli* DH5α to obtain clones. Plasmids from positive clones were then sequenced with T7 universal primers.

### High-throughput sequencing

The dsRNA products were mixed with primer1 (acgatgcaaagtctcgacaaatggtcgataccaattcaCCGCTCGAGCGGAGTTAACTTT) and dNTPs at 65 °C for 10 min and then cooled down to 24 °C for primer annealing. 10 × MMulV buffer, RNase inhibitor, and MMulV reverse transcriptase were added to the above system at 24 °C for 2 h to obtain cDNA, and then the enzymes were inactivated at 70 °C for 15 min. Subsequently, primer2 (cagataaactataattcctaatcgcgaggtggcactgcaaCGCGGATCCGCGGCTATGTGAGA) and high-fidelity DNA polymerase pfu-Phusion were added to perform PCR amplification. PCR products were then purified by agarose gel electrophoresis for high-throughput sequencing (Sangon Biotech).

### Substrate efficiency assay

RNA extension was performed by incubating 2 μM RdRp, 250 nM 13+23 nt RNA duplex, 50 μM substrate (which are NTPs, NDPs) in a 20 μl reaction system at 37 °C for 0, 10, 20, 30, 40, 50, 60 s. RdRp catalyzed reactions each containing three NTPs and one NDP (ADP GTP CTP UTP, ATP GDP CTP UTP, ATP GTP CDP UTP and ATP GTP CTP UDP) and 13+26 nt RNA duplex was performed for 1 min. The reactions were stopped with 2 × stop buffer. Grayscale analysis was then carried out using ImageJ and ilustrated by using OriginPro 8 (OriginLab, USA).

### RNA extension assays with MDP as substrate

A 13+26 nt RNA duplex was used in this assay (Fig.5e). The 5’ end of the RNA product strand was labeled with a 6-carboxyfluorescein (FAM) group, which allows us to monitor RNA elongation. RdRp can incorporate NHC monophosphate into RNA at the position where the nucleotide in the template is G or A. MDP and MTP were purchased from MedChemExpress.

For RNA extension RdRp at final concentration of 2 μM was incubated with 200 nM 13+26 nt RNA duplex and 50 μM ATP GTP UTP or CTP GTP UTP with or without 50 μM MTP/MDP in 20 μl reaction buffer for 1min at 37 °C. The reactions were stopped with 2 × stop buffer. 10 μl RNA product for each reaction was resolved on 20% denaturing polyacrylamide-urea gels and imaged with a Tanon-5200 Multi Fluorescence Imager to confirm the extension. Grayscale analysis was then carried out using ImageJ and ilustrated by using OriginPro 8 (OriginLab, USA).

### Incorporation mutation rate detection

RdRp at final concentration of 2 μM was incubated with 200 nM 13+29 nt RNA duplex and 50 μM ATP GTP UTP, 800 nM or 5 μM or 50 μM CTP and 50 μM MTP/MDP in 20 μl reaction buffer for 1min at 37 °C. Mutation rates were determined by High-throughput sequencing as mentioned above.

### Fluorescence assay for T7 RNA polymerase transcription

Transcription of T7 RNA polymerase was measured by a fluorescence assay which employs a pair of nucleic acid probes with fluorescent and quenching groups respectively to detect mRNA production. When the probes pair with the mRNA FAM and quenching group gets closer and the fluorescence produced by FAM will be quenched. The change of the fluorescent signal is proportional to the mRNA production.

## Data availability

All data generated or analyzed during this study are included in this published article (and its Supplementary Information files).

## Acknowledgements

The authors thank Yan Zhang at Zhejiang University for offering SARS-CoV-2 RdRp gene, Peihui Wang at Cheeloo College of Medicine, Shandong University for offering SARS-CoV RdRp gene, Haitao Yang at ShanghaiTech University for offering ZIKV NS5 gene, Wuxi Biortus Biosciences Co. Ltd for providing SARS-CoV-2 RdRp expressed in insect cells, Carina Muyao Gu at CBe-Learn School for helping with the English expression of the manuscript. This work was supported by Shandong Provincial Key Research and Development Program (2020CXGC011305).

## Author contributions

M.W. and L.G. conceived, designed the experiments and analyzed the data. M.W. performed functional assays of RdRp. M.W., C.W., and X.J. participated in expression, purification of the SARS-CoV-2 RdRp complex and other proteins. N.L., F.Z., H.D., S.W., M.C. helped with the experiments and participated in figure and manuscript preparation. M.W. and L.G. wrote the manuscript.

