## Supplementary Information files for "SARS-CoV-2 RdRp uses NDPs as a substrate and is able to incorporate NHC into RNA from diphosphate form molnupiravir": 0.8 μM CTP 50 μM MDP.docx

No mutations: TCCCTTT	56214-1269=54945 97.74%

Mutation: 1269

1: TCCCCTT	872

2: TCCCTTC	54

3: TCCTTT	68

4: TACCTTT	26

5: TCCCTCT	19

6: TCCATTT	16

7: TCCCGTT	12

8: TCTCTTT	11

9: TCCCTTA	137

10: TCACTTT	9

11: TCCCTAT	7

12: TCCCATT	8

13: ACCCTTT	6

14: TCCCTC	3

15: CCCCCTT	5

16: TGCCCTTT	3

17: TAAAAA	2

18: TCCCCTC	3

19: TGCCTTT	2

20: TCGCTTT	1

21: TCCCTGC	1

22: TCCCTGG	1

23: TCGCTT	1

24: TCCCCAT	1

25: TACCCTT	1

>Seq1_53688_56214_0.9551

GCTATGTGAGATTCCCTTTAAAGTTAACT

>Seq2_547_56214_0.0097

GCTATGTGAGATTCCCCTTAAAGTTAACT

>Seq3_148_56214_0.0026

GCTATGTGAGACTCCCTTTAAAGTTAACT

>Seq4_124_56214_0.0022

GCTATGTGAGATTCCCTTAAAGTTAACT

>Seq5_120_56214_0.0021

GCTATGTGAGATCCCCTTTAAAGTTAACT

>Seq6_88_56214_0.0016

GCTATGTGAGATCCCCTTAAAGTTAACT

>Seq7_87_56214_0.0015

GCTATGTGAGATTCCCCTTTAAAGTTAACT

>Seq8_63_56214_0.0011

GCTATGTGAGATTCCCTTTAAAGTTAACA

>Seq9_54_56214_0.0010

GCTATGTGAGATTCCCTTCAAAGTTAACT

>Seq10_42_56214_0.0007

GCTATGTGAGATTTCCCTTTAAAGTTAACT

>Seq11_40_56214_0.0007

GCTATGTGAGATTCCCTTTAAAGTTAACC

>Seq12_39_56214_0.0007

GCTATGTGAGATTCCCTTTAAAGTTAAC

>Seq13_39_56214_0.0007

GCTATGTGGGATTCCCTTTAAAGTTAACT

>Seq14_38_56214_0.0007

GCTATGTGAGGATTCCCTTTAAAGTTAACT

>Seq15_37_56214_0.0007

GCTATGTGAGATTCCCTTTAAGTTAACT

>Seq16_34_56214_0.0006

GCTATGCGAGATTCCCTTTAAAGTTAACT

>Seq17_31_56214_0.0006

CTATGTGAGATTCCCTTTAAAGTTAACT

>Seq18_30_56214_0.0005

GCTATGTGAGATTCCCTTTAAAAGTTAACT

>Seq19_29_56214_0.0005

GCTATGTGAGATTCCTTTAAAGTTAACT

>Seq20_28_56214_0.0005

GCTATGTGAGATTCCCTTTAAGGTTAACT

>Seq21_27_56214_0.0005

TCTATGTGAGATTCCCTTTAAAGTTAACT

>Seq22_27_56214_0.0005

GCTATGTGAGGTTCCCTTTAAAGTTAACT

>Seq23_26_56214_0.0005

GCTATGTGAGATTACCTTTAAAGTTAACT

>Seq24_25_56214_0.0004

GCTATGTGAGATTCCCTTTTAAAGTTAACT

>Seq25_25_56214_0.0004

GCTATGTGAGATTCCCTTTGAAGTTAACT

>Seq26_24_56214_0.0004

GCTATGTGAGATTTCCTTTAAAGTTAACT

>Seq27_24_56214_0.0004

GCTACGTGAGATTCCCTTTAAAGTTAACT

>Seq28_23_56214_0.0004

GCTATGTGAGATTCCCTTTAAAGTTAACTC

>Seq29_22_56214_0.0004

GCTATGTGAGATTCCCTTTAAAGCTAACT

>Seq30_21_56214_0.0004

GGCTATGTGAGATTCCCTTTAAAGTTAACT

>Seq31_20_56214_0.0004

GCTATGTGAGATTCCCTTTAAAGTTAGCT

>Seq32_20_56214_0.0004

GCTATGTGAGATTCCCTTTAAAGTTGACT

>Seq33_19_56214_0.0003

GCCATGTGAGATTCCCTTTAAAGTTAACT

>Seq34_19_56214_0.0003

GCTATGTGAGATTCCCTCTAAAGTTAACT

>Seq35_17_56214_0.0003

GTATGTGAGATTCCCTTTAAAGTTAACT

>Seq36_16_56214_0.0003

GCTATGTGAGATTCCCTTTAAAGTCAACT

>Seq37_16_56214_0.0003

GCTATGTGAGATTCCATTTAAAGTTAACT

>Seq38_14_56214_0.0002

GCTATGTGAGATTCCCTTTAGAGTTAACT

>Seq39_14_56214_0.0002

GTTATGTGAGATTCCCTTTAAAGTTAACT

>Seq40_13_56214_0.0002

GCTATGTGAGAATTCCCTTTAAAGTTAACT

>Seq41_13_56214_0.0002

GCTGTGTGAGATTCCCTTTAAAGTTAACT

>Seq42_12_56214_0.0002

GCTTGTGAGATTCCCTTTAAAGTTAACT

>Seq43_12_56214_0.0002

GCTATTTGAGATTCCCTTTAAAGTTAACT

>Seq44_12_56214_0.0002

GCTATGTGAGATTCCCTTTAAAATTAACT

>Seq45_12_56214_0.0002

GCTATGTGAGATTCCTTTTAAAGTTAACT

>Seq46_11_56214_0.0002

GCTATGTGAGATTCCCGTTAAAGTTAACT

>Seq47_10_56214_0.0002

GCTATGTGAGATTCTCTTTAAAGTTAACT

>Seq48_10_56214_0.0002

GCTATGTGAGATTCCCTTAAAAGTTAACT

>Seq49_10_56214_0.0002

GCTATGTGAGAATCCCCTTAAAGTTAACT

>Seq50_10_56214_0.0002

GCTATGAGAGATTCCCTTTAAAGTTAACT

>Seq51_10_56214_0.0002

GCTATGTGAGATTCCCTTTAAAGTTAAT

>Seq52_9_56214_0.0002

GCTATATGAGATTCCCTTTAAAGTTAACT

>Seq53_9_56214_0.0002

GCGATGTGAGATTCCCTTTAAAGTTAACT

>Seq54_9_56214_0.0002

GCATGTGAGATTCCCTTTAAAGTTAACT

>Seq55_9_56214_0.0002

ACTATGTGAGATTCCCTTTAAAGTTAACT

>Seq56_8_56214_0.0001

GCTATGTGATATTCCCTTTAAAGTTAACT

>Seq57_8_56214_0.0001

GCTATGTAAGATTCCCTTTAAAGTTAACT

>Seq58_8_56214_0.0001

GCTATGTGTGATTCCCTTTAAAGTTAACT

>Seq59_8_56214_0.0001

GCTATGTTAGATTCCCTTTAAAGTTAACT

>Seq60_8_56214_0.0001

GCTATGTGAGATTCACTTTAAAGTTAACT

>Seq61_8_56214_0.0001

GCTATGTGAGATTCCCTTTAAAGTTAATT

>Seq62_8_56214_0.0001

GCTATGTGAAATTCCCTTTAAAGTTAACT

>Seq63_8_56214_0.0001

GCAATGTGAGATTCCCTTTAAAGTTAACT

>Seq64_7_56214_0.0001

GCTATGTGAGATTCCCTTTAAAGTTATCT

>Seq65_7_56214_0.0001

GCTATGTGAGATTCCCTATAAAGTTAACT

>Seq66_7_56214_0.0001

GCTATGTGAGTTTCCCTTTAAAGTTAACT

>Seq67_7_56214_0.0001

GCTATGTGAGATCCCTTTAAAGTTAACT

>Seq68_7_56214_0.0001

GCTAAGTGAGATTCCCTTTAAAGTTAACT

>Seq69_6_56214_0.0001

GCTATGTGAGATTCCCATTAAAGTTAACT

>Seq70_5_56214_0.0001

GCTATGTGAGATTCCCTTTAAAGTTACT

>Seq71_5_56214_0.0001

GCTATGTGAGAATCCCTTTAAAGTTAACT

>Seq72_5_56214_0.0001

GCTATGTGAGATTCCCTTTAAAGATAACT

>Seq73_5_56214_0.0001

GCTATGTGAGATACCCTTTAAAGTTAACT

>Seq74_5_56214_0.0001

GCTATGTGAGATTCCCTTTTAAGTTAACT

>Seq75_5_56214_0.0001

GCTATGTGAGATTCCCTTTAAAGTTAACG

>Seq76_5_56214_0.0001

GCTTTGTGAGATTCCCTTTAAAGTTAACT

>Seq77_5_56214_0.0001

GCTATGTGAGATTCCCTTTAAAGTAACT

>Seq78_5_56214_0.0001

GCTATGTGAGATTCCCTTTAAAGTTAACCT

>Seq79_5_56214_0.0001

GCTATGTGAGATTCCCTTTAAAGTTAA

>Seq80_4_56214_0.0001

GCTATGTGAGATTCCCTTTAAAGTTAAAT

>Seq81_4_56214_0.0001

GCTATGTGAGATTCCCTTTAAAGTTAAGT

>Seq82_4_56214_0.0001

GCTATGTGAGATTCCCTTTAAAGTTAAACT

>Seq83_4_56214_0.0001

GCTATGTGCGATTCCCTTTAAAGTTAACT

>Seq84_4_56214_0.0001

GCTAGTGAGATTCCCTTTAAAGTTAACT

>Seq85_4_56214_0.0001

GCTATGGTGAGATTCCCTTTAAAGTTAACT

>Seq86_4_56214_0.0001

GCTATGTGAGATTCCCTTTAAAGTGAACT

>Seq87_4_56214_0.0001

GCTATGTGACATTCCCTTTAAAGTTAACT

>Seq88_4_56214_0.0001

GCTATGTGAGATTCCCTTTAAAGTTTACT

>Seq89_4_56214_0.0001

GCTAATGTGAGATTCCCTTTAAAGTTAACT

>Seq90_3_56214_0.0001

GCTATGTGAGATTCCCTTTATAGTTAACT

>Seq91_3_56214_0.0001

GGTATGTGAGATTCCCTTTAAAGTTAACT

>Seq92_3_56214_0.0001

GCTAGGTGAGATTCCCTTTAAAGTTAACT

>Seq93_3_56214_0.0001

GCCTATGTGAGATTCCCTTTAAAGTTAACT

>Seq94_3_56214_0.0001

GCTATGTGAGATTCCCTCAAAGTTAACT

>Seq95_3_56214_0.0001

GCTATGTGAGATTCCCTTTAAAGTTACCT

>Seq96_3_56214_0.0001

GCTATGGGAGATTCCCTTTAAAGTTAACT

>Seq97_3_56214_0.0001

GCTATGGAGATTCCCTTTAAAGTTAACT

>Seq98_3_56214_0.0001

GCTATGTGAGATTCCCTTTAAAGTTAACTT

>Seq99_3_56214_0.0001

GCTATGTGAGATTCCCCTTAAAGTTAACA

>Seq100_3_56214_0.0001

GCTATGTGAGATCCCCCTTAAAGTTAACT

>Seq101_2_56214_0.0000

GCTATGTGAGATTGCCCTTTAAAGTTAACT

>Seq102_2_56214_0.0000

GCTATGTGAGATTCCCTTTAATGTTAACT

>Seq103_2_56214_0.0000

GCTATGTGAGATTCCCTTTAAATTTAACT

>Seq104_2_56214_0.0000

GCTATGTGAGATTTCCCCTTAAAGTTAACT

>Seq105_2_56214_0.0000

GATGTGAGATTCCCTTTAAAGTTAACT

>Seq106_2_56214_0.0000

GCTATGTGAGATTAAAAAAAAGTTAACT

>Seq107_2_56214_0.0000

GCTATGTGAGATTCCCCTCAAAGTTAACT

>Seq108_2_56214_0.0000

GCTATGTGGAGATTCCCTTTAAAGTTAACT

>Seq109_2_56214_0.0000

GCTATGTGAAGATTCCCTTTAAAGTTAACT

>Seq110_2_56214_0.0000

GCTATGTGAGATTTTAAAGTTAACT

>Seq111_2_56214_0.0000

GCTATGTGAGATTCCCTTTAAAGTAAACT

>Seq112_2_56214_0.0000

GATATGTGAGATTCCCTTTAAAGTTAACT

>Seq113_2_56214_0.0000

GCTATGTGAGATTCCCTTTACAGTTAACT

>Seq114_2_56214_0.0000

GCTATGTGAGATTCCCTAAAGTTAACT

>Seq115_2_56214_0.0000

GCTATGTGAGATTGCCTTTAAAGTTAACT

>Seq116_2_56214_0.0000

GCTATGTGAGATTCCGTTTAAAGTTAACT

>Seq117_2_56214_0.0000

GCTATGTGAGACTCCCCTTAAAGTTAACT

>Seq118_1_56214_0.0000

GCTATGCGAGAATCCCCTTAAAGTTAACT

>Seq119_1_56214_0.0000

GCTATGTGAGATTCGCTTTAAAGTTAACT

>Seq120_1_56214_0.0000

GCTATGAGATTCCCTTTAAAGTTAACT

>Seq121_1_56214_0.0000

GCTATGTGAGATTCCCTTAAGGTTAACT

>Seq122_1_56214_0.0000

GCTATGTGAGATTCCTTTTAAGGTTAACT

>Seq123_1_56214_0.0000

GCTATGTGAGTTCCCTTTAAAGTTAACT

>Seq124_1_56214_0.0000

GCTATGTTGAGATTCCCTTTAAAGTTAACT

>Seq125_1_56214_0.0000

GCTTGTGAGATTCCCTTTAAAGTTAACTC

>Seq126_1_56214_0.0000

GCTATTTGAGATCCCCTTTAAAGTTAACT

>Seq127_1_56214_0.0000

GCTATGTGAGATTACCCTTTAAAGTTAACT

>Seq128_1_56214_0.0000

ACTATGTGAGGATTCCCTTTAAAGTTAACT

>Seq129_1_56214_0.0000

GCTATGTGAGATTCCCCTTTATAGTTAACT

>Seq130_1_56214_0.0000

GGTATGTGAGATTCCCTTTAAAGTTAACC

>Seq131_1_56214_0.0000

GCTATGTGAGATTCCCTTAAAGGTTAACT

>Seq132_1_56214_0.0000

GCTATGTGAGATTCCCTGCAAAGTTAACT

>Seq133_1_56214_0.0000

GCTATGTGATTCCCTTTAAAGTTAACT

>Seq134_1_56214_0.0000

GCTATGTGAGCTTCCCTTTAAAGTTAACT

>Seq135_1_56214_0.0000

GCTATGTGAGCTTCCCGTTAAAGTTAACT

>Seq136_1_56214_0.0000

GCTATGTGAGAGGCCCTTGAAAGGGCACT

>Seq137_1_56214_0.0000

GCTAGTGAGATTCCCCTTAAAGTTAACT

>Seq138_1_56214_0.0000

GCTATGTGAGATCCCCTCAAAGTTAACT

>Seq139_1_56214_0.0000

GCAGTGAGATTCCCTTTAAAGTTAACT

>Seq140_1_56214_0.0000

GCTATGTGAGACTCCCTTTAAAGTTAACC

>Seq141_1_56214_0.0000

GCTATGTGGGATTCCCCTTTAAAGTTAACT

>Seq142_1_56214_0.0000

GCTATGTGGATTCCCTTTAAAGTTAACT

>Seq143_1_56214_0.0000

GCTATGTGAGATTAAAGTTAACT

>Seq144_1_56214_0.0000

GCTGTGAGATTCCCTTTAAAGTTAACT

>Seq145_1_56214_0.0000

GCTATGCGAGACTCCCTTTAAAGTTAACT

>Seq146_1_56214_0.0000

GCTCTGTGAGATTCCCTTTAAAGTTAACT

>Seq147_1_56214_0.0000

GCTATGTGAGATTCCCTTTAAAGTTTACA

>Seq148_1_56214_0.0000

GCTATGGGAGATTCCCTGGAAAGTTCACT

>Seq149_1_56214_0.0000

GCTATTTGAGACTCCCTTTAAAGTTAACT

>Seq150_1_56214_0.0000

GCTATGTGAGATTTTCCTTTAAAGTTAACT

>Seq151_1_56214_0.0000

GCTATGCGAGATCCCCTTAAAGTTAACT

>Seq152_1_56214_0.0000

GCTATGTGAGATTCCCTTTAAAGTTCACT

>Seq153_1_56214_0.0000

GCTATGTGGAGGATTCCCTTTAAAGTTAACT

>Seq154_1_56214_0.0000

GCTATGTGAGATTCTTTAAAGTTAACT

>Seq155_1_56214_0.0000

GCTATGTGAGACTCCCATTAAAGTTAACT

>Seq156_1_56214_0.0000

GCTATGTGAGAGCCCCTTTAAAGTTAACT

>Seq157_1_56214_0.0000

GCTATGTGAGATTCCCTTTCAAGTTAACT

>Seq158_1_56214_0.0000

GCTATGTGAGATTCACTTTAAAATTAACT

>Seq159_1_56214_0.0000

GCTATTGTGAGATTCCCTTTAAAGTTAACT

>Seq160_1_56214_0.0000

GCTATGTGAGATTTAAAGTTAACT

>Seq161_1_56214_0.0000

GCTATGTGAGAATCCCCTTAAAGTTTACT

>Seq162_1_56214_0.0000

CGGCTATGTGAGATTCCCTTTAAAGTTAACT

>Seq163_1_56214_0.0000

GCTATGTTAGATTCCCTTAAAGTTAACT

>Seq164_1_56214_0.0000

GGCTATGTGAGATTCTCTTTAAAGTTAACT

>Seq165_1_56214_0.0000

GCTATGTGAGATTCGCTTAAAGTTAACT

>Seq166_1_56214_0.0000

GCTATGTGAGACCCCCTTTAAAGTTAACT

>Seq167_1_56214_0.0000

GCTATGTGAGATTCCCCTTGAAGTTAACT

>Seq168_1_56214_0.0000

GCTATGTGAGACTTCCCTTTAAAGTTAACT

>Seq169_1_56214_0.0000

GCTATGTGAGATTCCCCCTTAAAGTTAACT

>Seq170_1_56214_0.0000

GCTATGTGAGATCCCCTTAAAGTAAACT

>Seq171_1_56214_0.0000

GCTATGTGAGATGCCCTTTAAAGTTAACT

>Seq172_1_56214_0.0000

GCTATGTGAGATTTTTAAAGTTAACT

>Seq173_1_56214_0.0000

GCTATGTGAGATTTTTTAAAGTTAACT

>Seq174_1_56214_0.0000

GCTATGTGAGAGTCCCTTTAAAGTTAACT

>Seq175_1_56214_0.0000

GCTATCTGTGATTCCCTTTAAAGTTAACT

>Seq176_1_56214_0.0000

GCTACGCGAGATTCCCTTTAAAGTTAACT

>Seq177_1_56214_0.0000

GCTATTGAGATTCCCTTTAAAGTTAACT

>Seq178_1_56214_0.0000

GCTATGTGAATTCCCTTTAAAGTTAACT

>Seq179_1_56214_0.0000

GCTATGTGAGATCCCCTTTAATGTTAACT

>Seq180_1_56214_0.0000

GCTATGTGAGACTCCCTTTATAGTTAACT

>Seq181_1_56214_0.0000

GCTATGTGGGATTCCCCTTAAAAGTTAACT

>Seq182_1_56214_0.0000

GCTATGTGAGATTCCCTTTAAATTAACT

>Seq183_1_56214_0.0000

GCTATGTAGAGATTCCCTTTAAAGTTAACT

>Seq184_1_56214_0.0000

GATATGTGAGATTCCCTTTAGAGTTAACT

>Seq185_1_56214_0.0000

GCTATGAGAGATTCCCCTTAAAGTTAACT

>Seq186_1_56214_0.0000

GCTATGTGAGACTCCTTTAAAGTTAACT

>Seq187_1_56214_0.0000

GCTATGTGAGATTCCCTTTAAAGTTTAACT

>Seq188_1_56214_0.0000

GCTATGAGAGATTCCCATTAAAGTTAACT

>Seq189_1_56214_0.0000

GCTACGTGAGATCCCCTTTAAAGTTAACT

>Seq190_1_56214_0.0000

GCTATGTGAGATTCCCCATAAAGTTAACT

>Seq191_1_56214_0.0000

GCTTATGTGAGATTCCCTTTAAAGTTAACT

>Seq192_1_56214_0.0000

GCTATGTGAGATTACCCTTAAAGTTAACT

>Seq193_1_56214_0.0000

GCTGTGTGGGATTCCCTTTAAAGTTAACT
