## Supplementary Information files for "SARS-CoV-2 RdRp uses NDPs as a substrate and is able to incorporate NHC into RNA from diphosphate form molnupiravir": 0.8 μM CTP 50 μM MTP.docx

No mutations: TCCCTTT	59129-1298=57831 97.80%

Mutations: 1298

1: TCCCCTT	887

2: TCCCTTC	58

3: TTCCTTT	77

4: TCCATTT	29

5: TCCCTCT	19

6: TCCCGTT	16

7: TACCTTT	15

8: TCCCATT	13

9: TCACTTT	13

10: TCTCTTT	11

11: TCCCTTA	124

12: TCCCCTC	7

13: TCCCTAT	4

14: ACCCTTT	5

15: CCCCCTT	6

16: TGCCTTT	3

17: TGCCCTTT	3

18: TCGCCTT	2

19: TCCCAGTT	2

20: TCCCTGT	2

21: TCCCTCAG	1

22: TCGCTTT	1

23: TCCCCCCCC	1

24: TTCCTCT	1

25: TCCCCCC	1

26: TCCCCT	1

27: TAAAAAA	1

28: TCACCTT	1

>Seq1_56559_59129_0.9565

GCTATGTGAGATTCCCTTTAAAGTTAACT

>Seq2_559_59129_0.0095

GCTATGTGAGATTCCCCTTAAAGTTAACT

>Seq3_153_59129_0.0026

GCTATGTGAGACTCCCTTTAAAGTTAACT

>Seq4_112_59129_0.0019

GCTATGTGAGATTCCCTTAAAGTTAACT

>Seq5_111_59129_0.0019

GCTATGTGAGATCCCCTTTAAAGTTAACT

>Seq6_97_59129_0.0016

GCTATGTGAGATTCCCCTTTAAAGTTAACT

>Seq7_94_59129_0.0016

GCTATGTGAGATCCCCTTAAAGTTAACT

>Seq8_61_59129_0.0010

GCTATGTGAGATTCCCTTTAAAGTTAACA

>Seq9_56_59129_0.0009

GCTATGTGAGATTCCCTTCAAAGTTAACT

>Seq10_51_59129_0.0009

GCTATGCGAGATTCCCTTTAAAGTTAACT

>Seq11_43_59129_0.0007

GCTATGTGAGATTTCCCTTTAAAGTTAACT

>Seq12_43_59129_0.0007

GCTATGTGAGATTCCCTTTAAAGTTAAC

>Seq13_41_59129_0.0007

GCTATGTGGGATTCCCTTTAAAGTTAACT

>Seq14_39_59129_0.0007

GCTATGTGAGATTCCCTTTAAAGTTAACC

>Seq15_35_59129_0.0006

GCTATGTGAGATTCCCTTTAAGTTAACT

>Seq16_35_59129_0.0006

GCTATGTGAGGATTCCCTTTAAAGTTAACT

>Seq17_33_59129_0.0006

GCTATGTGAGATTCCCTTTTAAAGTTAACT

>Seq18_33_59129_0.0006

GCTATGTGAGATTTCCTTTAAAGTTAACT

>Seq19_28_59129_0.0005

GCCATGTGAGATTCCCTTTAAAGTTAACT

>Seq20_28_59129_0.0005

GCTATGTGAGATTCCATTTAAAGTTAACT

>Seq21_27_59129_0.0005

GCTATGTGAGGTTCCCTTTAAAGTTAACT

>Seq22_26_59129_0.0004

GCTATGTGAGATTCCTTTAAAGTTAACT

>Seq23_26_59129_0.0004

CTATGTGAGATTCCCTTTAAAGTTAACT

>Seq24_26_59129_0.0004

GCTACGTGAGATTCCCTTTAAAGTTAACT

>Seq25_23_59129_0.0004

GCTATGTGAGAATTCCCTTTAAAGTTAACT

>Seq26_23_59129_0.0004

GCTATGTGAGATTCCCTTTAAGGTTAACT

>Seq27_23_59129_0.0004

GCTATGTGAGATTCCCTTTAAAGCTAACT

>Seq28_22_59129_0.0004

GCTATGTGAGATTCCCTTTAAAGTCAACT

>Seq29_21_59129_0.0004

GCTATGTGAGATTCCCTTTAAAGTTAGCT

>Seq30_19_59129_0.0003

GCTATGTGAGATTCCCTTTAAAAGTTAACT

>Seq31_18_59129_0.0003

GCTATGTGTGATTCCCTTTAAAGTTAACT

>Seq32_17_59129_0.0003

GCTATGTGAGATTCCCTCTAAAGTTAACT

>Seq33_17_59129_0.0003

GCTATGTGAGATTCCCTTTAAAGTTGACT

>Seq34_16_59129_0.0003

GCTATGTAAGATTCCCTTTAAAGTTAACT

>Seq35_16_59129_0.0003

GGCTATGTGAGATTCCCTTTAAAGTTAACT

>Seq36_16_59129_0.0003

TCTATGTGAGATTCCCTTTAAAGTTAACT

>Seq37_16_59129_0.0003

GCTATGTGAGATTCCCGTTAAAGTTAACT

>Seq38_15_59129_0.0003

GCTATGTGAGATTACCTTTAAAGTTAACT

>Seq39_15_59129_0.0003

GCTATGTGAGATTCCTTTTAAAGTTAACT

>Seq40_14_59129_0.0002

GCTATGTGAGATTCCCTTTAGAGTTAACT

>Seq41_14_59129_0.0002

GCATGTGAGATTCCCTTTAAAGTTAACT

>Seq42_13_59129_0.0002

GCTATGTGAGATTCCCTTTAAAGTTAACTC

>Seq43_13_59129_0.0002

GCTATGTGAGATTCCCATTAAAGTTAACT

>Seq44_13_59129_0.0002

GCTATGTGAGATTCACTTTAAAGTTAACT

>Seq45_13_59129_0.0002

ACTATGTGAGATTCCCTTTAAAGTTAACT

>Seq46_12_59129_0.0002

GCTATGTGAGATTCCCTTTAAAGTTACT

>Seq47_12_59129_0.0002

GCTATGTGAGAATCCCCTTAAAGTTAACT

>Seq48_11_59129_0.0002

GCTATGTGATATTCCCTTTAAAGTTAACT

>Seq49_11_59129_0.0002

GCTATGTGAGATTCCCTTTGAAGTTAACT

>Seq50_11_59129_0.0002

GCTATGTGAGATTCCCTTTTAAGTTAACT

>Seq51_10_59129_0.0002

GCTTGTGAGATTCCCTTTAAAGTTAACT

>Seq52_10_59129_0.0002

GCTATGTGAGATTCTCTTTAAAGTTAACT

>Seq53_10_59129_0.0002

GCTATGTGAGATTCCCTTAAAAGTTAACT

>Seq54_10_59129_0.0002

GCTATGTGAGATTCCCTTTAAAATTAACT

>Seq55_9_59129_0.0002

GTTATGTGAGATTCCCTTTAAAGTTAACT

>Seq56_9_59129_0.0002

GCTATGTGCGATTCCCTTTAAAGTTAACT

>Seq57_9_59129_0.0002

GCTATATGAGATTCCCTTTAAAGTTAACT

>Seq58_9_59129_0.0002

GCTATGTGAGATCCCTTTAAAGTTAACT

>Seq59_9_59129_0.0002

GCTATGTGAGATTCCCTTTAAAGTTAACCT

>Seq60_9_59129_0.0002

GCTATGAGAGATTCCCTTTAAAGTTAACT

>Seq61_9_59129_0.0002

GCTATGTGAGATTCCCTTTAAAGTTAAT

>Seq62_8_59129_0.0001

GTATGTGAGATTCCCTTTAAAGTTAACT

>Seq63_8_59129_0.0001

GCTGTGTGAGATTCCCTTTAAAGTTAACT

>Seq64_8_59129_0.0001

GCGATGTGAGATTCCCTTTAAAGTTAACT

>Seq65_7_59129_0.0001

GCTATGTGAGATTCCCTTTAAAGTTAAAT

>Seq66_7_59129_0.0001

GCTATTTGAGATTCCCTTTAAAGTTAACT

>Seq67_7_59129_0.0001

GCTATGTGAGATTCCCTTTAAAGTTAATT

>Seq68_7_59129_0.0001

GCTATGTGAAATTCCCTTTAAAGTTAACT

>Seq69_6_59129_0.0001

GCTATGTGAGAATCCCTTTAAAGTTAACT

>Seq70_6_59129_0.0001

GCTATGTTAGATTCCCTTTAAAGTTAACT

>Seq71_6_59129_0.0001

GCTAGTGAGATTCCCTTTAAAGTTAACT

>Seq72_6_59129_0.0001

GCTATGTGAGATTCCCTTTAAAGTTAA

>Seq73_6_59129_0.0001

GCAATGTGAGATTCCCTTTAAAGTTAACT

>Seq74_5_59129_0.0001

GCTATGTGAGTTTCCCTTTAAAGTTAACT

>Seq75_5_59129_0.0001

GCTATGTGAGATTCCCCTCAAAGTTAACT

>Seq76_5_59129_0.0001

GCTATGTGAGATTCCCTTTAAAGTTAACG

>Seq77_5_59129_0.0001

GCTTTGTGAGATTCCCTTTAAAGTTAACT

>Seq78_5_59129_0.0001

GCCTATGTGAGATTCCCTTTAAAGTTAACT

>Seq79_5_59129_0.0001

GCTATGTGAGATTCCCTTTAAAGTAAACT

>Seq80_5_59129_0.0001

GCTATGTGACATTCCCTTTAAAGTTAACT

>Seq81_5_59129_0.0001

GCTATGTGAGATTCCCTTTAAAGTTTACT

>Seq82_5_59129_0.0001

GCTAATGTGAGATTCCCTTTAAAGTTAACT

>Seq83_5_59129_0.0001

GCTATGTGAGATTCCCTTTAAAGTTAACTT

>Seq84_4_59129_0.0001

GCTATGTGAGATTCCCTTTAATGTTAACT

>Seq85_4_59129_0.0001

GCTATGTGAGATTCCCTTTAAAGTTATCT

>Seq86_4_59129_0.0001

GCTATGTGAGATTCCCTATAAAGTTAACT

>Seq87_4_59129_0.0001

CCTATGTGAGATTCCCTTTAAAGTTAACT

>Seq88_4_59129_0.0001

GCTATGTGAGATTCCCTTTAAAGATAACT

>Seq89_4_59129_0.0001

GGTATGTGAGATTCCCTTTAAAGTTAACT

>Seq90_4_59129_0.0001

GCTATGTGAGATTCCCTTTAAAGTTAAACT

>Seq91_4_59129_0.0001

GCTATGTGAGATACCCTTTAAAGTTAACT

>Seq92_4_59129_0.0001

GCTATGTGAAGATTCCCTTTAAAGTTAACT

>Seq93_4_59129_0.0001

GCTAAGTGAGATTCCCTTTAAAGTTAACT

>Seq94_4_59129_0.0001

GCTATGTGAGAGTCCCTTTAAAGTTAACT

>Seq95_4_59129_0.0001

GCTATGGGAGATTCCCTTTAAAGTTAACT

>Seq96_3_59129_0.0001

GCTATCTGAGATTCCCTTTAAAGTTAACT

>Seq97_3_59129_0.0001

GCTATGTGAGATTCCCTTTAAAGTAACT

>Seq98_3_59129_0.0001

GCTATGTGAGATTTTTAAAGTTAACT

>Seq99_3_59129_0.0001

GCTATGTGAGATCCCCCTTAAAGTTAACT

>Seq100_3_59129_0.0001

GCTATGTGAGATTGCCTTTAAAGTTAACT

>Seq101_2_59129_0.0000

GCTATGTGAGATTGCCCTTTAAAGTTAACT

>Seq102_2_59129_0.0000

GCTATGAGAGATTCCCTTTAAAGTAACT

>Seq103_2_59129_0.0000

GCTATGTGAGATTCCCTTTATAGTTAACT

>Seq104_2_59129_0.0000

GCTATGTGAGATCCCCCTTTAAAGTTAACT

>Seq105_2_59129_0.0000

GCTATGTGAGATCCCCTCAAAGTTAACT

>Seq106_2_59129_0.0000

GCTAGGTGAGATTCCCTTTAAAGTTAACT

>Seq107_2_59129_0.0000

GATGTGAGATTCCCTTTAAAGTTAACT

>Seq108_2_59129_0.0000

GCTGTGAGATTCCCTTTAAAGTTAACT

>Seq109_2_59129_0.0000

GCTATGTGAGATTCGCCTTAAAGTTAACT

>Seq110_2_59129_0.0000

GCTATGTGGAGATTCCCTTTAAAGTTAACT

>Seq111_2_59129_0.0000

GCTATGTGAGATTCTTTAAAGTTAACT

>Seq112_2_59129_0.0000

GCTATGCGAGATTCCCCTTAAAGTTAACT

>Seq113_2_59129_0.0000

GCTATTGTGAGATTCCCTTTAAAGTTAACT

>Seq114_2_59129_0.0000

GCTATGTGAGATTCCCAGTTAAAGTTAACT

>Seq115_2_59129_0.0000

GCTATGTGAGATTCCCTTTACAGTTAACT

>Seq116_2_59129_0.0000

GCTATGTGAGATTCCCTGTAAAGTTAACT

>Seq117_2_59129_0.0000

GCTATGTGAGATTCCCTAAAGTTAACT

>Seq118_2_59129_0.0000

GCTATGTGAGATTCCCTTTAAAGTTTAACT

>Seq119_2_59129_0.0000

GCTATGTGAGACTCCCTTTAAAGTTAAC

>Seq120_1_59129_0.0000

GCTATGTGAGGATCCCCTTAAAGTTAACT

>Seq121_1_59129_0.0000

GCTATGTGAGATTCCCTCAGAAGTTAACT

>Seq122_1_59129_0.0000

GCATGTGAGATTCCCTTTGAAGTTAACT

>Seq123_1_59129_0.0000

GCTATGTGAGATTCGCTTTAAAGTTAACT

>Seq124_1_59129_0.0000

GCTATGTGAGATTCCCTTTAAAGTTA

>Seq125_1_59129_0.0000

CTATGTGAGTTTCCCTTTAAAGTTAACT

>Seq126_1_59129_0.0000

GCTATGTGAGATTACCCTTTAAAGTTAACT

>Seq127_1_59129_0.0000

GCTATGTGAGATTCCTTTAAAGTTAGCT

>Seq128_1_59129_0.0000

GCTATGTGGGATTCTCTTTAAAGTTAACT

>Seq129_1_59129_0.0000

GCTATGTGAGACTCCCTCTAAAGTTAACT

>Seq130_1_59129_0.0000

CTATGTGAGACTCCCTTTAAAGTTAACT

>Seq131_1_59129_0.0000

GCTATGTGAGATTCCCTTTAAAGTTAAGT

>Seq132_1_59129_0.0000

GCTATGTGAGATTCCCCTTAAAGTTAACC

>Seq133_1_59129_0.0000

GCTATGTGAGATTCCCTTTAAAGCTTAACT

>Seq134_1_59129_0.0000

GCTGTGTGAGATTCCCCTTAAAGTTAACT

>Seq135_1_59129_0.0000

GCTATGTGAGATTCCCTTTAAATTTAACT

>Seq136_1_59129_0.0000

TTTATGTGAGATTCCCTTTAAAGTTAACT

>Seq137_1_59129_0.0000

CTATGTGAGATTCCCTTCAAAGTTAACT

>Seq138_1_59129_0.0000

GCTATGTGAGATTCCCCTTAAAATTAACT

>Seq139_1_59129_0.0000

GGCTATGTGAGATTCCCTCTAAAGTTAACT

>Seq140_1_59129_0.0000

GCTATGTGAGATTCCCCCCCCAAAGTTAACT

>Seq141_1_59129_0.0000

GCTATGCGAGATTCCCTTTAAAGTTAACA

>Seq142_1_59129_0.0000

GCTATGTGAGACTCCCTTAAAGTTAACT

>Seq143_1_59129_0.0000

GGCTATGTGAGATTCCCCTTAAAGTTAACT

>Seq144_1_59129_0.0000

GCTATGTGAGAATTCCCTTAAAGTTAACT

>Seq145_1_59129_0.0000

GCTATGTGAGATTTCCTCTAAAGTTAACT

>Seq146_1_59129_0.0000

GCTATGTGAGATCTTTAAAGTTAACT

>Seq147_1_59129_0.0000

GCTATGTGAGATTTCCCCTTAAAGTTAACT

>Seq148_1_59129_0.0000

GCTATGTGAGATCCCCTTTAAAGTTAACA

>Seq149_1_59129_0.0000

GCTATGTGAGATTCCCCCCAAAGTTAACT

>Seq150_1_59129_0.0000

TCTATGTGAGATTCCCCTTAAAGTTAACT

>Seq151_1_59129_0.0000

GCTATGTGGATTCCCTTTAAAGTTAACT

>Seq152_1_59129_0.0000

GCTATGTGAGATTCCCCTAAAGTTAACT

>Seq153_1_59129_0.0000

GCTATGTGAGATTACCCCTTTAAAGTTAACT

>Seq154_1_59129_0.0000

GCTATGCGAGACTCCCTTTAAAGTTAACT

>Seq155_1_59129_0.0000

GCTCTGTGAGATTCCCTTTAAAGTTAACT

>Seq156_1_59129_0.0000

GCTATGTGAGATTAAAAAAAAAGTTAACT

>Seq157_1_59129_0.0000

GCTATGTGAGATTCCATTTGAAAGTTAACT

>Seq158_1_59129_0.0000

GCCATGTGAGATTCCCTTTAAAGTCAACT

>Seq159_1_59129_0.0000

GCTATGGTGAGATTCCCTTTAAAGTTAACT

>Seq160_1_59129_0.0000

GCTATGTGAGATTCACCTTAAAGTTAACT

>Seq161_1_59129_0.0000

GCTATGTGAGATTTAAAGTTAACT

>Seq162_1_59129_0.0000

GCGTGAGATTCCCTTTAAAGTTAACT

>Seq163_1_59129_0.0000

GCTATGTGAGATTTTAAAGTTAACT

>Seq164_1_59129_0.0000

GCTTGTGAGATTCCCCTTAAAGTTAACT

>Seq165_1_59129_0.0000

GCTATGTCAGATTCCCTTTAAAGTTAACT

>Seq166_1_59129_0.0000

GCTATGTGAGATTCCCCCTTAAAGTTAACT

>Seq167_1_59129_0.0000

GCTATGTGAGATGCCCTTTAAAGTTAACT

>Seq168_1_59129_0.0000

GATATGTGAGATTCCCTTTAAAGTTAACT

>Seq169_1_59129_0.0000

GCTATGTGAGATTCCCTTTAAAGGTAACT

>Seq170_1_59129_0.0000

GCTATGTGAGAATCCCCTTTAAGTTAACT

>Seq171_1_59129_0.0000

GCTATGTGAGACTCCCTTTAAAGTTAACA

>Seq172_1_59129_0.0000

GCTACGTGAGATCCCCTTAAAGTTAACT

>Seq173_1_59129_0.0000

GCTATTGAGATTCCCTTTAAAGTTAACT

>Seq174_1_59129_0.0000

GCTATGTGAGATTCCCTTTAAAGTTACCT

>Seq175_1_59129_0.0000

GCTACGTGAGATTCCCTTCAAAGTTAACT

>Seq176_1_59129_0.0000

GCTATGGAGATTCCCTTTAAAGTTAACT

>Seq177_1_59129_0.0000

GCTATGTGAGATCCCCCTCAAAGTTAACT

>Seq178_1_59129_0.0000

TCTATGTGAGATTTCCTTTAAAGTTAACT

>Seq179_1_59129_0.0000

TCTAGTGAGATTCCCTTTAAAGTTAACT

>Seq180_1_59129_0.0000

GCTATGTAGATTCCCTTTAAAGTTAACT

>Seq181_1_59129_0.0000

GCTTATGTGAGATTCCCTTTAAAGTTAACT

>Seq182_1_59129_0.0000

GCTATGTGAGGATTCCTTTAAAGTTAACT
