## Supplementary Information files for "SARS-CoV-2 RdRp uses NDPs as a substrate and is able to incorporate NHC into RNA from diphosphate form molnupiravir": 5 μM CTP 50 μM MDP.docx

No mutations 70906-1463=69443 97.94%

mutations	1463

1: TCCCCTT	1007

2: TCCCTT	215

3: TCCCTCT	34

4: TCCTTT	47

5: TCCATTT	22

6: TACCTTT	19

7: TTCCTTT	21

8: TCCCGTT	16

9: TCACTTT	12

10: TCCCATT	10

11: TCTCTTT	8

12: TCCCTAT	9

13: TCCCCTC	9

14: TCTTT	4

15: TCGCTTT	3

16: TCCGTTT	3

17: TCCCTC	4

18: TCCCCCTT	6

19: TCCCTGT	3

20: TACCCTT	5

21: CCCCGC	1

22: TAACCTTT	1

23: ACCCCTT	1

24: TAAAAAA	1

25: TCCCTGC	1

26: TGCCCTTT	1

>Seq1_63292_66026_0.9586

GCTATGTGAGATTCCCTTTAAAGTTAACT

>Seq2_564_66026_0.0085

GCTATGTGAGATTCCCCTTAAAGTTAACT

>Seq3_209_66026_0.0032

GCTATGTGAGACTCCCTTTAAAGTTAACT

>Seq4_136_66026_0.0021

GCTATGTGAGATTCCCTTAAAGTTAACT

>Seq5_129_66026_0.0020

GCTATGTGAGATCCCCTTTAAAGTTAACT

>Seq6_126_66026_0.0019

GCTATGTGAGATTCCCCTTTAAAGTTAACT

>Seq7_73_66026_0.0011

GCTATGTGAGATTCCCTTTAAAGTTAACA

>Seq8_71_66026_0.0011

GCTATGTGAGATCCCCTTAAAGTTAACT

>Seq9_58_66026_0.0009

GCTATGTGAGATTCCCTTCAAAGTTAACT

>Seq10_54_66026_0.0008

GCTATGCGAGATTCCCTTTAAAGTTAACT

>Seq11_46_66026_0.0007

GCTATGTGGGATTCCCTTTAAAGTTAACT

>Seq12_39_66026_0.0006

GCTATGTGAGATTCCCTTTTAAAGTTAACT

>Seq13_35_66026_0.0005

CTATGTGAGATTCCCTTTAAAGTTAACT

>Seq14_33_66026_0.0005

GCTATGTGAGATTCCCTTTAAGTTAACT

>Seq15_32_66026_0.0005

GCTATGTGAGATTCCCTTTAAAGTTAAC

>Seq16_31_66026_0.0005

GCTATGTGAGATTCCCTCTAAAGTTAACT

>Seq17_30_66026_0.0005

GCTATGTGAGATTCCCTTTAAAGTTAACC

>Seq18_30_66026_0.0005

GCTATGTGAGATTCCCTTTAAAGCTAACT

>Seq19_30_66026_0.0005

GCTATGTGAGATTTCCCTTTAAAGTTAACT

>Seq20_30_66026_0.0005

GCTATGTGAGATTCCCTTTAAAAGTTAACT

>Seq21_28_66026_0.0004

GGCTATGTGAGATTCCCTTTAAAGTTAACT

>Seq22_28_66026_0.0004

GCTATGTGAGATTCCTTTAAAGTTAACT

>Seq23_28_66026_0.0004

GCCATGTGAGATTCCCTTTAAAGTTAACT

>Seq24_25_66026_0.0004

TCTATGTGAGATTCCCTTTAAAGTTAACT

>Seq25_25_66026_0.0004

GCTATGTGAGGATTCCCTTTAAAGTTAACT

>Seq26_25_66026_0.0004

GCTACGTGAGATTCCCTTTAAAGTTAACT

>Seq27_25_66026_0.0004

GCTATGTGAGATTCCCTTTAAAGTTGACT

>Seq28_23_66026_0.0003

GCTATGTGAGATTCCCTTTAAAGTTAGCT

>Seq29_22_66026_0.0003

GCTATGTGAGGTTCCCTTTAAAGTTAACT

>Seq30_21_66026_0.0003

GCTATGTGAGAATTCCCTTTAAAGTTAACT

>Seq31_21_66026_0.0003

GCTATGTGAGATTTCCTTTAAAGTTAACT

>Seq32_21_66026_0.0003

GCTATGTGAGATTCCATTTAAAGTTAACT

>Seq33_20_66026_0.0003

GCTATGTGAGATTCCCTTTAAAGTCAACT

>Seq34_19_66026_0.0003

GCTATGTGAGATTCCCTTTAAAGTTAACTC

>Seq35_18_66026_0.0003

GCTATGTGAGAATCCCCTTAAAGTTAACT

>Seq36_17_66026_0.0003

GCTATGTGAGATTACCTTTAAAGTTAACT

>Seq37_17_66026_0.0003

GCTATGTGTGATTCCCTTTAAAGTTAACT

>Seq38_17_66026_0.0003

GCTATGTGAGATTCCCTTTAAGGTTAACT

>Seq39_17_66026_0.0003

GCTATATGAGATTCCCTTTAAAGTTAACT

>Seq40_16_66026_0.0002

GCTATGTAAGATTCCCTTTAAAGTTAACT

>Seq41_15_66026_0.0002

GTTATGTGAGATTCCCTTTAAAGTTAACT

>Seq42_15_66026_0.0002

GCTATGTGAGATTCCCGTTAAAGTTAACT

>Seq43_14_66026_0.0002

GCTATGTGAGATTCCCTTTAGAGTTAACT

>Seq44_14_66026_0.0002

GCATGTGAGATTCCCTTTAAAGTTAACT

>Seq45_14_66026_0.0002

ACTATGTGAGATTCCCTTTAAAGTTAACT

>Seq46_12_66026_0.0002

GCTATGTTAGATTCCCTTTAAAGTTAACT

>Seq47_12_66026_0.0002

GCTATGTGAGATTCCCTTTAAAATTAACT

>Seq48_11_66026_0.0002

GCTATGTGAGATTCACTTTAAAGTTAACT

>Seq49_11_66026_0.0002

GCTATGTGAGATTCCCTTTAAAGTTAACCT

>Seq50_11_66026_0.0002

GCGATGTGAGATTCCCTTTAAAGTTAACT

>Seq51_11_66026_0.0002

GCTATGTGAGATTCCTTTTAAAGTTAACT

>Seq52_10_66026_0.0002

GCTATGTGAGATTCCCATTAAAGTTAACT

>Seq53_10_66026_0.0002

GCTATGTGAGATACCCTTTAAAGTTAACT

>Seq54_10_66026_0.0002

GCTATGTGAGATTCCCTTTGAAGTTAACT

>Seq55_10_66026_0.0002

GCTATGTGAGATTCCCTTTAAAGTTAAT

>Seq56_9_66026_0.0001

GTATGTGAGATTCCCTTTAAAGTTAACT

>Seq57_9_66026_0.0001

GCTTGTGAGATTCCCTTTAAAGTTAACT

>Seq58_9_66026_0.0001

GCTATGTGAGTTTCCCTTTAAAGTTAACT

>Seq59_9_66026_0.0001

GCTATTTGAGATTCCCTTTAAAGTTAACT

>Seq60_9_66026_0.0001

GCTATGTGAGATTCCCTTTAAAGTTAATT

>Seq61_8_66026_0.0001

GCTATGTGAGATTCCCTATAAAGTTAACT

>Seq62_8_66026_0.0001

GCTATGTGAGAATCCCTTTAAAGTTAACT

>Seq63_8_66026_0.0001

GCTATGTGAGATTCTCTTTAAAGTTAACT

>Seq64_8_66026_0.0001

GCTATGTGAGAGTCCCTTTAAAGTTAACT

>Seq65_7_66026_0.0001

GCTATGTGCGATTCCCTTTAAAGTTAACT

>Seq66_7_66026_0.0001

GCTATGTGAGATTCCCTTTAAAGTTAACG

>Seq67_7_66026_0.0001

GCTATGTGAGATTCCCTTTAAAGTTAA

>Seq68_7_66026_0.0001

GCTATGTGAAATTCCCTTTAAAGTTAACT

>Seq69_6_66026_0.0001

GCTATGTGAGATTCCCTTTAAAGTTATCT

>Seq70_6_66026_0.0001

GCTATGTGAGATTCCCTTTAAAGTTACT

>Seq71_6_66026_0.0001

GCTATGTGAGATTCCCTTTAAAGATAACT

>Seq72_6_66026_0.0001

GCTATGTGAGATTCCCCTCAAAGTTAACT

>Seq73_6_66026_0.0001

GCTAGTGAGATTCCCTTTAAAGTTAACT

>Seq74_6_66026_0.0001

GCTATGTGAGATCCCTTTAAAGTTAACT

>Seq75_6_66026_0.0001

GCTATGTGAGATTCCCTTTAAAGTAAACT

>Seq76_6_66026_0.0001

GCTAAGTGAGATTCCCTTTAAAGTTAACT

>Seq77_6_66026_0.0001

GCTATGTGAGATTCCCTTAAAAGTTAACT

>Seq78_6_66026_0.0001

GCTATGAGAGATTCCCTTTAAAGTTAACT

>Seq79_5_66026_0.0001

GCTATGTGATATTCCCTTTAAAGTTAACT

>Seq80_5_66026_0.0001

GCTATGTGAGATTCCCTTTTAAGTTAACT

>Seq81_5_66026_0.0001

GCTATGTGAGATTCCCTTTAAAGGTAACT

>Seq82_5_66026_0.0001

GCTATGTGAGATTCCCTTTAAAGTTAACTT

>Seq83_4_66026_0.0001

GCTATGTTGAGATTCCCTTTAAAGTTAACT

>Seq84_4_66026_0.0001

GCTCTGTGAGATTCCCTTTAAAGTTAACT

>Seq85_4_66026_0.0001

GCTATGTGGAGATTCCCTTTAAAGTTAACT

>Seq86_4_66026_0.0001

GCTATGTGAGATTCTTTAAAGTTAACT

>Seq87_4_66026_0.0001

GCTTTGTGAGATTCCCTTTAAAGTTAACT

>Seq88_4_66026_0.0001

GCCTATGTGAGATTCCCTTTAAAGTTAACT

>Seq89_4_66026_0.0001

GCTATGTGAGATTCCCTTTAAAGTTTACT

>Seq90_4_66026_0.0001

GCTATGTAGATTCCCTTTAAAGTTAACT

>Seq91_4_66026_0.0001

GCTATGTGAGACTCCCCTTAAAGTTAACT

>Seq92_3_66026_0.0000

GCTATGTGAGATTCCCTTTAAAGTTAAAT

>Seq93_3_66026_0.0000

GCTATGTGAGATTCGCTTTAAAGTTAACT

>Seq94_3_66026_0.0000

GCTGTGTGAGATTCCCTTTAAAGTTAACT

>Seq95_3_66026_0.0000

GCTAGGTGAGATTCCCTTTAAAGTTAACT

>Seq96_3_66026_0.0000

GCTATGTGAAGATTCCCTTTAAAGTTAACT

>Seq97_3_66026_0.0000

GCTATGTGACATTCCCTTTAAAGTTAACT

>Seq98_3_66026_0.0000

GCTATGTCAGATTCCCTTTAAAGTTAACT

>Seq99_3_66026_0.0000

GATATGTGAGATTCCCTTTAAAGTTAACT

>Seq100_3_66026_0.0000

GCTATGGAGATTCCCTTTAAAGTTAACT

>Seq101_3_66026_0.0000

GCAATGTGAGATTCCCTTTAAAGTTAACT

>Seq102_3_66026_0.0000

GCTATGTGAGATTCCCTTGAAAGTTAACT

>Seq103_3_66026_0.0000

GCTATGTGAGATTCCGTTTAAAGTTAACT

>Seq104_2_66026_0.0000

GCTATGTGAGATTCCCTTTATAGTTAACT

>Seq105_2_66026_0.0000

CATGTGAGATTCCCTTTAAAGTTAACT

>Seq106_2_66026_0.0000

GCTATGTGAGATTCCCCTTAAAGTTAAC

>Seq107_2_66026_0.0000

GCTATGTGAGATTAAAGTTAACT

>Seq108_2_66026_0.0000

GCTATGTGAGATTTTCCTTTAAAGTTAACT

>Seq109_2_66026_0.0000

GCTATGTGAGATTCCCTTTAAAGTGAACT

>Seq110_2_66026_0.0000

GCTATGTGAGATTCCCTTTAAAGTAACT

>Seq111_2_66026_0.0000

GCTATGTGAGATTCCCCCTTAAAGTTAACT

>Seq112_2_66026_0.0000

GCTATGTGAGATTCCCTCAAAGTTAACT

>Seq113_2_66026_0.0000

GCTATGTGAGATTCCCTTTACAGTTAACT

>Seq114_2_66026_0.0000

GCTATGTGAGATTCCCTGTAAAGTTAACT

>Seq115_2_66026_0.0000

GCTATGGGAGATTCCCTTTAAAGTTAACT

>Seq116_2_66026_0.0000

GCTATGTGAATTCCCTTTAAAGTTAACT

>Seq117_2_66026_0.0000

GCTAATGTGAGATTCCCTTTAAAGTTAACT

>Seq118_2_66026_0.0000

GCTATGTGAGATCCCCCTTAAAGTTAACT

>Seq119_1_66026_0.0000

GTATGTGAGAATCCCCTTAAAGTTAACT

>Seq120_1_66026_0.0000

GCTATATGAGATTCCCTTTAAAGTTAGCT

>Seq121_1_66026_0.0000

GCTATGTGAGATTGCCCTTTAAAGTTAACT

>Seq122_1_66026_0.0000

GCTATGTGAGGATCCCCTTAAAGTTAACT

>Seq123_1_66026_0.0000

GCTATGTGAGATTCCCTGAAAGTTAACT

>Seq124_1_66026_0.0000

GCTATGTGAGATTTCCCTTAAAGTTAACT

>Seq125_1_66026_0.0000

GCTATGTGAGATTCCCTTTAATGTTAACT

>Seq126_1_66026_0.0000

GCTATGTGAGATTCCCTTTAACGTTAACT

>Seq127_1_66026_0.0000

GCTAGGTGAGATTCCCCTTAAAGTTAACT

>Seq128_1_66026_0.0000

GCTATGTAAGACTCCCTTTAAAGTTAACT

>Seq129_1_66026_0.0000

GCTATGTGAGATTCCCTTAAGGTTAACT

>Seq130_1_66026_0.0000

GCTATGTGAGATTCCTAAAGTTAACT

>Seq131_1_66026_0.0000

GCTATGTGAGATTCCCTTTATGTTAACT

>Seq132_1_66026_0.0000

GCTATGTGAGATTCCCTGTTAAAGTTAACT

>Seq133_1_66026_0.0000

CCTATGTGAGATTCCCTTTAAAGTTAACT

>Seq134_1_66026_0.0000

GCTTGTGAGAATTCCCTTTAAAGTTAACT

>Seq135_1_66026_0.0000

GCTATGAGAGATCCCCCTTAAAGTTAACT

>Seq136_1_66026_0.0000

GCTATCTGAGATTCCCTTTAAAGTTAACT

>Seq137_1_66026_0.0000

GCTATGTGAGATTACCCTTTAAAGTTAACT

>Seq138_1_66026_0.0000

GCTATGTGAGATTCCCCTTAAAGTTAACTT

>Seq139_1_66026_0.0000

GCTATGTGAGCTTCCCTTTAAAGTTACCT

>Seq140_1_66026_0.0000

GGTATGTGAGATTCCCTTTAAAGTTAACT

>Seq141_1_66026_0.0000

GCTATGTGAGATTCCCTTTAAAGTTAAGT

>Seq142_1_66026_0.0000

GCTATGCGAGATTCCCTTTAAAGATAACT

>Seq143_1_66026_0.0000

GCTATGTGAGATTCCCTGCAAAGTTAACT

>Seq144_1_66026_0.0000

GCTATGTGAGATACCCTTTAAGGTTAACT

>Seq145_1_66026_0.0000

GCTATGTGAGATACCCCTTAAAGTTAACT

>Seq146_1_66026_0.0000

GCTATGTGAGATTCCCTTTAAATTTAACT

>Seq147_1_66026_0.0000

GCTATGTGTGATTCCCCTTTAAAGTTAACT

>Seq148_1_66026_0.0000

ATGTGAGATTCCCTTTAAAGTTAACT

>Seq149_1_66026_0.0000

GCTATGTGAGATTCCCTTTAAAGTTAAACT

>Seq150_1_66026_0.0000

GCTATGTGAGCTTCCCTTTAAAGTTAACT

>Seq151_1_66026_0.0000

GGCTATGTGAGATTCCCTTAAAAGTTAACT

>Seq152_1_66026_0.0000

GCTATGTGAGATTCCCCTTTAAGTTAACT

>Seq153_1_66026_0.0000

GCTATGTGAGATTCACTTTAAAAGTTAACT

>Seq154_1_66026_0.0000

GTTATGTGAGATTCCTTTAAAGTTAACT

>Seq155_1_66026_0.0000

GCTATGTGAGATTCCCTTTAAAGTCAACA

>Seq156_1_66026_0.0000

GCTATGTGAGATTCCCCTTAAAGTTAGCT

>Seq157_1_66026_0.0000

GCTATGTGAGATTCCCTTTAAAGTACT

>Seq158_1_66026_0.0000

GGCTATGTGAGATTCCCCTTTAAAGTTAACT

>Seq159_1_66026_0.0000

GCTATGTGGATTCCCTTTAAAGTTAACT

>Seq160_1_66026_0.0000

GTGAGATTTCCCTTTAAAGTTAACT

>Seq161_1_66026_0.0000

GCTATGTGAGATTAAAAAAAAAGTTAACT

>Seq162_1_66026_0.0000

GCTATGTGAGATCCCCTTTATAGTTAACT

>Seq163_1_66026_0.0000

GCTATGGTGAGATTCCCTTTAAAGTTAACT

>Seq164_1_66026_0.0000

GCTATGTGAGATTCCCTTTAAAGTTCACT

>Seq165_1_66026_0.0000

GCTATGTGAGATTCCCTTTCAAGTTAACT

>Seq166_1_66026_0.0000

GCTATGTGAGATCCCCTTAAAAGTTAACT

>Seq167_1_66026_0.0000

GCTAGTGAGATTCCCTTTAAAGTCAACT

>Seq168_1_66026_0.0000

GCGTGAGATTCCCTTTAAAGTTAACT

>Seq169_1_66026_0.0000

GCTATGTGAGATTTTAAAGTTAACT

>Seq170_1_66026_0.0000

GCTATGTGAGATTCCCCCTTTAAAGTTAACT

>Seq171_1_66026_0.0000

CTATGTGAGATCCCCTTTAAAGTTAACT

>Seq172_1_66026_0.0000

GCTATGTGAGACTCCCTCTTAAAGTTAACT

>Seq173_1_66026_0.0000

GCTATGTGAGATCCACTTAAAGTTAACT

>Seq174_1_66026_0.0000

GCATGTGAGATTCCCTTTAAAGTTAATT

>Seq175_1_66026_0.0000

GCTATGTGAGATTTTTAAAGTTAACT

>Seq176_1_66026_0.0000

GCTATGTGGGATTTCCCTTTAAAGTTAACT

>Seq177_1_66026_0.0000

GGTATGTGAGATTCCCCTTAAAGTTAACT

>Seq178_1_66026_0.0000

GCTATGTGAGATACCCTTTAGAGTTAACT

>Seq179_1_66026_0.0000

GCTATGTGAGGATTCCCTTAAAGTTAACT

>Seq180_1_66026_0.0000

GCTATGTGAGATTCCCTTTAAAGTTACCT

>Seq181_1_66026_0.0000

GCTATGTGAGATCCCCGCAAAGTTAACT

>Seq182_1_66026_0.0000

GCTATGTGAGATTCCCCTTAAAGGTAACT

>Seq183_1_66026_0.0000

GCTATGTGAGATTCCCCTTTAAAGATAACT

>Seq184_1_66026_0.0000

GCTATGAGAGATTCCCCTTAAAGTTAACT

>Seq185_1_66026_0.0000

CTATGTGAGATTCCCTTTAAAGTCAACT

>Seq186_1_66026_0.0000

GCTATGTGAGATTCCCTTGAAGTTAACT

>Seq187_1_66026_0.0000

GCTATGTGGACATTCCCTTTAAAGTTAACT

>Seq188_1_66026_0.0000

GCTATGTGAGATCCCCTCTAAAGTTAACT

>Seq189_1_66026_0.0000

GCTATGTGAGATTCCCTTTAAAGTTTAACT

>Seq190_1_66026_0.0000

GCTATGTGAGATTCCCTTTAAAAGTTAACA

>Seq191_1_66026_0.0000

GCTATGTGAGATTAACCTTTAAAGTTAACT

>Seq192_1_66026_0.0000

GCTATGTGAGATTCCCCTTAAAGTTAACA

>Seq193_1_66026_0.0000

GCTTATGTGAGATTCCCTTTAAAGTTAACT

>Seq194_1_66026_0.0000

GCTATGTGAGATTACCCTTAAAGTTAACT

>Seq195_1_66026_0.0000

GCTATGAGAGATTCCCTTTAAGTTAACT

>Seq196_1_66026_0.0000

TATGTGAGATTCCCTTTAAAGTTAACT
