## Supplementary Information files for "SARS-CoV-2 RdRp uses NDPs as a substrate and is able to incorporate NHC into RNA from diphosphate form molnupiravir": 5 μM CTP 50 μM MTP.docx

no mutations: TCCCTTT 59588-1202=58386 97.98%

mutations	1202

1: TCCCCTT	772

2: TCCCTT	205

3: TTCCTTT	51

4: TCCCTCT	33

5: TACCTTT	26

6: TCTCTTT	17

7: TCCCATT	15

8: TCACTTT	13

9: TCCCGTT	9

10: TCCATTT	9

11: TCCCCTC	9

12: CCCCCTT	6

13: TCCCTAT	4

14: TCCCTGT	4

15: GCCCTTT	6

16: TGCCTTT	5

17: TCCCTGC	2

18: TCCCAGTT 2

19: TCCCTC	1

20: TAAAAAC 1

21: ACCCTTT	4

22: TTCTTTT	1

23: TAACCTTT	1

24: CCCCCTC	1

25: CCCGATGC	1

26: GTCCCTGG	1

27: TCGCTTT	1

28: TCCGTTT	1

29: TCCCTG	1

>Seq1_52851_55135_0.9586

GCTATGTGAGATTCCCTTTAAAGTTAACT

>Seq2_384_55135_0.0070

GCTATGTGAGATTCCCCTTAAAGTTAACT

>Seq3_137_55135_0.0025

GCTATGTGAGACTCCCTTTAAAGTTAACT

>Seq4_132_55135_0.0024

GCTATGTGAGATTCCCTTAAAGTTAACT

>Seq5_117_55135_0.0021

GCTATGTGAGATCCCCTTTAAAGTTAACT

>Seq6_107_55135_0.0019

GCTATGTGAGATTCCCCTTTAAAGTTAACT

>Seq7_69_55135_0.0013

GCTATGTGAGATCCCCTTAAAGTTAACT

>Seq8_57_55135_0.0010

GCTATGTGAGATTCCCTTTAAAGTTAACA

>Seq9_51_55135_0.0009

GCTATGTGAGATTCCCTTCAAAGTTAACT

>Seq10_40_55135_0.0007

GCTATGTGGGATTCCCTTTAAAGTTAACT

>Seq11_38_55135_0.0007

GCTATGTGAGATTCCCTTTTAAAGTTAACT

>Seq12_38_55135_0.0007

GCTATGTGAGATTCCCTTTAAAGTTAAC

>Seq13_37_55135_0.0007

GCTATGTGAGATTTCCCTTTAAAGTTAACT

>Seq14_33_55135_0.0006

CTATGTGAGATTCCCTTTAAAGTTAACT

>Seq15_32_55135_0.0006

GCTATGTGAGGATTCCCTTTAAAGTTAACT

>Seq16_32_55135_0.0006

GCTATGCGAGATTCCCTTTAAAGTTAACT

>Seq17_29_55135_0.0005

GCTATGTGAGATTCCCTTTAAAGTTAACC

>Seq18_29_55135_0.0005

GCTATGTGAGATTCCCTTTAAGGTTAACT

>Seq19_26_55135_0.0005

GCTATGTGAGATTCCCTTTAAGTTAACT

>Seq20_26_55135_0.0005

GCTATGTGAGATTCCCTTTAAAAGTTAACT

>Seq21_25_55135_0.0005

GCTACGTGAGATTCCCTTTAAAGTTAACT

>Seq22_24_55135_0.0004

TCTATGTGAGATTCCCTTTAAAGTTAACT

>Seq23_24_55135_0.0004

GCTATGTGAGATTTCCTTTAAAGTTAACT

>Seq24_23_55135_0.0004

GCTATGTGAGGTTCCCTTTAAAGTTAACT

>Seq25_23_55135_0.0004

GCTATGTGAGATTCCCTCTAAAGTTAACT

>Seq26_21_55135_0.0004

GGCTATGTGAGATTCCCTTTAAAGTTAACT

>Seq27_21_55135_0.0004

GCTATGTGAGATTCCCTTTGAAGTTAACT

>Seq28_19_55135_0.0003

GCTATGTGAGATTACCTTTAAAGTTAACT

>Seq29_19_55135_0.0003

GCTATGTGAGATTCCTTTAAAGTTAACT

>Seq30_18_55135_0.0003

GCCATGTGAGATTCCCTTTAAAGTTAACT

>Seq31_18_55135_0.0003

GCTATGTGAGATTCCCTTTAAAGTCAACT

>Seq32_17_55135_0.0003

GCTATGTGAGATTCCCTTTAAAGCTAACT

>Seq33_15_55135_0.0003

GCTATGTGAGATTCCCTTTAGAGTTAACT

>Seq34_15_55135_0.0003

GCTATGTGAGAATTCCCTTTAAAGTTAACT

>Seq35_15_55135_0.0003

GCTATGTGAGATTCCCTTTAAAGTTAGCT

>Seq36_15_55135_0.0003

GCTATGTGAGATTCTCTTTAAAGTTAACT

>Seq37_14_55135_0.0003

GCTATGTGAGATTCCCATTAAAGTTAACT

>Seq38_14_55135_0.0003

GCTATGTGAGATTCACTTTAAAGTTAACT

>Seq39_13_55135_0.0002

GCTATATGAGATTCCCTTTAAAGTTAACT

>Seq40_13_55135_0.0002

ACTATGTGAGATTCCCTTTAAAGTTAACT

>Seq41_13_55135_0.0002

GCTATGTGAGATTCCTTTTAAAGTTAACT

>Seq42_12_55135_0.0002

GCTATGTAAGATTCCCTTTAAAGTTAACT

>Seq43_12_55135_0.0002

GCTAGTGAGATTCCCTTTAAAGTTAACT

>Seq44_12_55135_0.0002

GCTATTTGAGATTCCCTTTAAAGTTAACT

>Seq45_11_55135_0.0002

GCTTGTGAGATTCCCTTTAAAGTTAACT

>Seq46_11_55135_0.0002

GCTATGTGAGATTCCCTTTAAAGTTAACTC

>Seq47_11_55135_0.0002

GCTATGTGTGATTCCCTTTAAAGTTAACT

>Seq48_10_55135_0.0002

GCTATGTGATATTCCCTTTAAAGTTAACT

>Seq49_10_55135_0.0002

GCTGTGTGAGATTCCCTTTAAAGTTAACT

>Seq50_10_55135_0.0002

GCTATGTTAGATTCCCTTTAAAGTTAACT

>Seq51_10_55135_0.0002

GCTATGTGAGAATCCCCTTAAAGTTAACT

>Seq52_10_55135_0.0002

GCGATGTGAGATTCCCTTTAAAGTTAACT

>Seq53_9_55135_0.0002

GTATGTGAGATTCCCTTTAAAGTTAACT

>Seq54_8_55135_0.0001

GTTATGTGAGATTCCCTTTAAAGTTAACT

>Seq55_8_55135_0.0001

GCTATGTGAGATTCCCTTTAAAGTTAATT

>Seq56_8_55135_0.0001

GCTATGTGAGATTCCCTTAAAAGTTAACT

>Seq57_8_55135_0.0001

GCTATGTGAGATTCCCGTTAAAGTTAACT

>Seq58_8_55135_0.0001

GCATGTGAGATTCCCTTTAAAGTTAACT

>Seq59_8_55135_0.0001

GCTATGTGAAATTCCCTTTAAAGTTAACT

>Seq60_8_55135_0.0001

GCTATGAGAGATTCCCTTTAAAGTTAACT

>Seq61_8_55135_0.0001

GCTATGTGAGATTCCCTTTAAAGTTGACT

>Seq62_7_55135_0.0001

GCTATGTGAGATTCCCTTTAAAGATAACT

>Seq63_7_55135_0.0001

GCTATGTGAGTTTCCCTTTAAAGTTAACT

>Seq64_7_55135_0.0001

GCTATGTGCGATTCCCTTTAAAGTTAACT

>Seq65_7_55135_0.0001

GCTATGTGAGATTCCCTTTTAAGTTAACT

>Seq66_7_55135_0.0001

GCTATGTGAGATTCCATTTAAAGTTAACT

>Seq67_7_55135_0.0001

GCTATGTGAGATTCCCTTTAAAGTTAAT

>Seq68_7_55135_0.0001

GCTATGTGAGATTCCCTTTAAAATTAACT

>Seq69_6_55135_0.0001

GCTATGTGAGATTCCCCTCAAAGTTAACT

>Seq70_6_55135_0.0001

GCTATGTGAGATTCCCTTTAAAGTTAACG

>Seq71_6_55135_0.0001

GCAATGTGAGATTCCCTTTAAAGTTAACT

>Seq72_5_55135_0.0001

GCTATGTGAGATTCCCTATAAAGTTAACT

>Seq73_5_55135_0.0001

GCTATGTGAGAATCCCTTTAAAGTTAACT

>Seq74_5_55135_0.0001

GCTATGTGAAGATTCCCTTTAAAGTTAACT

>Seq75_5_55135_0.0001

GCTATGTGAGATTCCCTTTAAAGTTAACCT

>Seq76_5_55135_0.0001

GATATGTGAGATTCCCTTTAAAGTTAACT

>Seq77_5_55135_0.0001

GCTATGTGAGATTCCCTGTAAAGTTAACT

>Seq78_5_55135_0.0001

GCTAATGTGAGATTCCCTTTAAAGTTAACT

>Seq79_4_55135_0.0001

GCTATGTGAGATTCCCTTTATAGTTAACT

>Seq80_4_55135_0.0001

GCTATGTGAGATTCCCTTTAAAGTTATCT

>Seq81_4_55135_0.0001

GCTATGTGAGATTCCCTTTAAAGTTAAGT

>Seq82_4_55135_0.0001

GCTTTGTGAGATTCCCTTTAAAGTTAACT

>Seq83_4_55135_0.0001

GCTATGTGAGATTCCCTTTAAAGTTTACT

>Seq84_4_55135_0.0001

GCTATGTGAGATTCCCTTTAAAGGTAACT

>Seq85_4_55135_0.0001

GCTATGTGAGATTCCCTTTAAAGTTAACTT

>Seq86_4_55135_0.0001

GCTATGTGAGATCCCCCTTAAAGTTAACT

>Seq87_4_55135_0.0001

GCTATGTGAGATTGCCTTTAAAGTTAACT

>Seq88_3_55135_0.0001

GCTATGTGAGATTGCCCTTTAAAGTTAACT

>Seq89_3_55135_0.0001

GGTATGTGAGATTCCCTTTAAAGTTAACT

>Seq90_3_55135_0.0001

GCTATGTGAGATTCCCTGCAAAGTTAACT

>Seq91_3_55135_0.0001

GCTATGTGAGATTCCCTTTAAAGTTAAACT

>Seq92_3_55135_0.0001

GCTAGGTGAGATTCCCTTTAAAGTTAACT

>Seq93_3_55135_0.0001

GCTATGTGAGATTTAAAGTTAACT

>Seq94_3_55135_0.0001

GCTATGTGAGATTCCCTTTAAAGTAACT

>Seq95_3_55135_0.0001

GCTAAGTGAGATTCCCTTTAAAGTTAACT

>Seq96_3_55135_0.0001

GCTATGTGAGATGCCCTTTAAAGTTAACT

>Seq97_3_55135_0.0001

GCTATGTGAGATTCCCTTTAAAGTTAA

>Seq98_3_55135_0.0001

GCTATGTGAGATTCCCTTTAAAGTTACCT

>Seq99_3_55135_0.0001

GCTATGTGAATTCCCTTTAAAGTTAACT

>Seq100_2_55135_0.0000

GCTATGTGAGATTCCCTTTAAAGTTAAAT

>Seq101_2_55135_0.0000

GCTATGTGAGATTCCCTTTAATGTTAACT

>Seq102_2_55135_0.0000

GCTATGTGAGATTCCCTTTAAAGTTACT

>Seq103_2_55135_0.0000

CCTATGTGAGATTCCCTTTAAAGTTAACT

>Seq104_2_55135_0.0000

GCTATCTGAGATTCCCTTTAAAGTTAACT

>Seq105_2_55135_0.0000

GCTATGTTGAGATTCCCTTTAAAGTTAACT

>Seq106_2_55135_0.0000

GCTATGTGAGATTCCCTTTAAATTTAACT

>Seq107_2_55135_0.0000

GCTATGTGAGATTTCCCCTTAAAGTTAACT

>Seq108_2_55135_0.0000

GCTATGTGAGATTAAAGTTAACT

>Seq109_2_55135_0.0000

GCTATGTGGAGATTCCCTTTAAAGTTAACT

>Seq110_2_55135_0.0000

GCTATGGTGAGATTCCCTTTAAAGTTAACT

>Seq111_2_55135_0.0000

GCTATGTGAGATTCCCTTTAAAGTGAACT

>Seq112_2_55135_0.0000

GCTATGTGAGATCCCTTTAAAGTTAACT

>Seq113_2_55135_0.0000

GCGTGAGATTCCCTTTAAAGTTAACT

>Seq114_2_55135_0.0000

GCTATGTGAGATTTTAAAGTTAACT

>Seq115_2_55135_0.0000

GCTATGTGAGATTCCCAGTTAAAGTTAACT

>Seq116_2_55135_0.0000

GCTATGTGACATTCCCTTTAAAGTTAACT

>Seq117_2_55135_0.0000

GCTATGTGAGACCCCCTTTAAAGTTAACT

>Seq118_2_55135_0.0000

GCTATGTGAGATTCTTAAAGTTAACT

>Seq119_2_55135_0.0000

GCTATGCGAGATTCCCTTAAAGTTAACT

>Seq120_1_55135_0.0000

GCTATGTGAGACTCCCTTTAAAGTTAAT

>Seq121_1_55135_0.0000

GCTATGTGAGATTCCCTGAAAGTTAACT

>Seq122_1_55135_0.0000

TCTATGTGAGATTCCTTTAAAGTTAACT

>Seq123_1_55135_0.0000

GCTATGTGAGACTCCCTTCAAAGTTAACT

>Seq124_1_55135_0.0000

GCTATGTGAGATTCCTTTTAAAGTTAACA

>Seq125_1_55135_0.0000

GCTATGTGAGATTCCCTTTAACGTTAACT

>Seq126_1_55135_0.0000

GCTATGTGAGATTCGCTTTAAAGTTAACT

>Seq127_1_55135_0.0000

GCTACGTGAGGTTCCCTTTAAAGTTAACT

>Seq128_1_55135_0.0000

GCTATGTGAGACTTCCTTTAAAGTTAACT

>Seq129_1_55135_0.0000

GCTATGTGAGATTCCCCTTAAAGTCAACT

>Seq130_1_55135_0.0000

GCTAGTGAGATGCCCTTTAAAGTTAACT

>Seq131_1_55135_0.0000

GCTATGAGATTCCCTTTAAAGTTAACT

>Seq132_1_55135_0.0000

GCTATGTGAGATTCCTAAAGTTAACT

>Seq133_1_55135_0.0000

GCTATGTGAGAATCCCCTCAAAGTTAACT

>Seq134_1_55135_0.0000

GCTATGTGAGATTAAAAAAGTTAACT

>Seq135_1_55135_0.0000

GCTATGTGAGATTCCCTTTAATAGTTAACT

>Seq136_1_55135_0.0000

GCTATGCGAGATTCCTTTTAAAGTTAACT

>Seq137_1_55135_0.0000

CCTATGTGAGATTCCCCTTAAAGTTAACT

>Seq138_1_55135_0.0000

GCTATGTTAGATTCCCCTTAAAGTTAACT

>Seq139_1_55135_0.0000

GCTATGTGGGATTCCCTTTAAAGTTACT

>Seq140_1_55135_0.0000

GCTATGTGACATTCCCCTTAAAGTTAACT

>Seq141_1_55135_0.0000

GCTATGTGAGATTCCCCTTTAAAGTTAAC

>Seq142_1_55135_0.0000

GCTATGTGAGAGTCCCCTTAAAGTTAACT

>Seq143_1_55135_0.0000

GCTGTGTGAGATTCCCCTTAAAGTTAACT

>Seq144_1_55135_0.0000

GCTATGTGAGATTCCCTGCAAGGTTAACT

>Seq145_1_55135_0.0000

GCTATGTGAGATTAAGTTAACT

>Seq146_1_55135_0.0000

GCTATGTGAGATTCAAAGTTAACT

>Seq147_1_55135_0.0000

GCTATGTGAGATTCCCTTAAAGTTAACA

>Seq148_1_55135_0.0000

GCTATGTGATATCCCCCTTAAAGTTAACT

>Seq149_1_55135_0.0000

CTATGTGAGATTCCCTTTAAAGCTAACT

>Seq150_1_55135_0.0000

GCTATGTGAGATTTTCCCTTTAAAGTTAACT

>Seq151_1_55135_0.0000

GATGTGAGATTCCCTTTAAAGTTAACT

>Seq152_1_55135_0.0000

GCCATATGAGATTCCCTTTAAAGTTAACT

>Seq153_1_55135_0.0000

GCTATGTGAGATTCCCCTTAAAGTTAGCT

>Seq154_1_55135_0.0000

GCTATGTGAGATTAAAAACAAAGTTAACT

>Seq155_1_55135_0.0000

GCTATGTGAGATTCCCCTAAAGTTAACT

>Seq156_1_55135_0.0000

GCTATGTGAGATACCCTTTAAAGTTAACT

>Seq157_1_55135_0.0000

GCTGTGAGATTCCCTTTAAAGTTAACT

>Seq158_1_55135_0.0000

GTATGTGAGATTTCTTTTAAAGTTAACT

>Seq159_1_55135_0.0000

GCTTGTGAGATTACCTTTAAAGTTAACT

>Seq160_1_55135_0.0000

GCTATGCGAGGTTCCCTTTAAAGTTAACT

>Seq161_1_55135_0.0000

GCTATGTGAGATTCCCTTTAAAGTTCACT

>Seq162_1_55135_0.0000

GCTATGTGAGATTCCCTTTAAATGTTAACT

>Seq163_1_55135_0.0000

GCTATGTGAGACTCCCTTTAAGTTAACT

>Seq164_1_55135_0.0000

GCTATGTGAGATTCCCTTTCAAGTTAACT

>Seq165_1_55135_0.0000

GCTATTGTGAGATTCCCTTTAAAGTTAACT

>Seq166_1_55135_0.0000

CTATGTGAGGTTCCCTTTAAAGTTAACT

>Seq167_1_55135_0.0000

GCTATGTGAGATTCCCTTTTAAAGTTAAACT

>Seq168_1_55135_0.0000

GCTAGTGAGATTCCCCTTTAAAGTTAACT

>Seq169_1_55135_0.0000

GCTATGTGAGGTTCCCCTTAAAGTTAACT

>Seq170_1_55135_0.0000

GCCTATGTGAGATTCCCTTTAAAGTTAACT

>Seq171_1_55135_0.0000

GATATGTGAGACTCCCTTTAAAGTTAACT

>Seq172_1_55135_0.0000

GCTATGTGAGATTCCCATTTAAAGTTAACT

>Seq173_1_55135_0.0000

GCTATGTGAGATTCCCTTTAAAGTAAACT

>Seq174_1_55135_0.0000

GTATGTGAGATTCCCCTTTAAAGTTAACT

>Seq175_1_55135_0.0000

GCTATGTGAGATTCCCTTTAAAGTTAACAC

>Seq176_1_55135_0.0000

GCTATGTGAGACTCCCCTCAAAGTTAACT

>Seq177_1_55135_0.0000

GCTATGTGAGACTTCCCTTTAAAGTTAACT

>Seq178_1_55135_0.0000

GCTATGTGAGATTCCATTTAAAGTAACT

>Seq179_1_55135_0.0000

GCTATGTGAGATTTTTAAAGTTAACT

>Seq180_1_55135_0.0000

GCTACGCGAGATTCCCTTTAAAGTTAACT

>Seq181_1_55135_0.0000

GCTATGTGAGATTCCCCTTTAAAGTTAACTC

>Seq182_1_55135_0.0000

GCTATGTGAGATTCCCTTTACAGTTAACT

>Seq183_1_55135_0.0000

GCTATGTGAGATCCCGATGCAAAGTTAACT

>Seq184_1_55135_0.0000

GCTATGGGAGATTCCCTTTAAAGTTAACT

>Seq185_1_55135_0.0000

ACTATGTGAGATTTCCCTTTAAAGTTAACT

>Seq186_1_55135_0.0000

GCTATGGAGATTCCCTTTAAAGTTAACT

>Seq187_1_55135_0.0000

GCTATGTGAGATTCCCATTTGTAAAGTTAACT

>Seq188_1_55135_0.0000

GCTATGTGAGATTCCCCTTTAAAGTTAGCT

>Seq189_1_55135_0.0000

GCTACGTGAGATTCCCCTTAAAGTTAACT

>Seq190_1_55135_0.0000

GCTATGTGAGATCCCCCTCAAAGTTAACT

>Seq191_1_55135_0.0000

GCTATGTGAGATTCCCTCAAAGTTAAC

>Seq192_1_55135_0.0000

GCTATGTGAGATTCCCTTTAAAGCTAA

>Seq193_1_55135_0.0000

GCTATGTGAGGATTCCCTTTGAAGTTAACT

>Seq194_1_55135_0.0000

GCTATGCGAGATTCCCCTTTAAAGTTAACT

>Seq195_1_55135_0.0000

GCTATGTGAGATTCACTTTAAAGTTAACA

>Seq196_1_55135_0.0000

GCCATGTGAGATTCCCCTTAAAGTTAACT

>Seq197_1_55135_0.0000

GCTATGTGAGATTCCCCTTAAAGTTAACA

>Seq198_1_55135_0.0000

GCTATGTGAGATTAACCTTTAAAGTTAACT

>Seq199_1_55135_0.0000

GCTTATGTGAGATTCCCTTTAAAGTTAACT

>Seq200_1_55135_0.0000

GCTATGTGCGATTCCCTTTGAAGTTAACT

>Seq201_1_55135_0.0000

GCTATGTGGGACTCCCTTTAAAGTTAACT

>Seq202_1_55135_0.0000

GCTATGTGAGATTCCCCTTAAAGTTACT

>Seq203_1_55135_0.0000

TATGTGAGATTCCCTTTAAAGTTAACT

>Seq204_1_55135_0.0000

GCTATGTGAGATTCCGTTTAAAGTTAACT
