## Supplementary Information files for "SARS-CoV-2 RdRp uses NDPs as a substrate and is able to incorporate NHC into RNA from diphosphate form molnupiravir": 13+23nt RNA NDP.docx

>Seq1_61830_63294_0.9769

GCTATGTGAGATTAAAGTTAACT

>Seq2_255_63294_0.0040

GCTATGTGAGATCAAAGTTAACT

>Seq3_104_63294_0.0016

GCTATGTGAGGATTAAAGTTAACT

>Seq4_62_63294_0.0010

GCTATGTGAGATTAAAAGTTAACT

>Seq5_52_63294_0.0008

CTATGTGAGATTAAAGTTAACT

>Seq6_43_63294_0.0007

GCTATGTGAGATTAAAGTTAAC

>Seq7_41_63294_0.0006

GCTATGTGAGATTTAAAGTTAACT

>Seq8_31_63294_0.0005

GCTATGTGGGATTAAAGTTAACT

>Seq9_31_63294_0.0005

GCTATGTGAGATTAAAGTTAACA

>Seq10_30_63294_0.0005

GCTATGTGAGATGAAAGTTAACT

>Seq11_30_63294_0.0005

GCTATGTGAGAATTAAAGTTAACT

>Seq12_28_63294_0.0004

GCTATGTGAGATTAAGTTAACT

>Seq13_26_63294_0.0004

GCTATGTGAGATTAAAGTTAAT

>Seq14_25_63294_0.0004

GCTATGTGAGATTAAAGTTACT

>Seq15_22_63294_0.0003

GTATGTGAGATTAAAGTTAACT

>Seq16_21_63294_0.0003

GCATGTGAGATTAAAGTTAACT

>Seq17_21_63294_0.0003

GCTACGTGAGATTAAAGTTAACT

>Seq18_20_63294_0.0003

GCTATGCGAGATTAAAGTTAACT

>Seq19_19_63294_0.0003

GGCTATGTGAGATTAAAGTTAACT

>Seq20_18_63294_0.0003

GCTATGTGAGATTAGAGTTAACT

>Seq21_18_63294_0.0003

GCTATGTGAGATTGAAGTTAACT

>Seq22_17_63294_0.0003

GTTATGTGAGATTAAAGTTAACT

>Seq23_17_63294_0.0003

GCTATGTGGAGATTAAAGTTAACT

>Seq24_17_63294_0.0003

GCTATGTGAGATTAAAGTTGACT

>Seq25_17_63294_0.0003

GCTATATGAGATTAAAGTTAACT

>Seq26_17_63294_0.0003

GCTATGTGAGATTAAAGTTAACC

>Seq27_15_63294_0.0002

TCTATGTGAGATTAAAGTTAACT

>Seq28_15_63294_0.0002

GCTATGTGAGATTAAAGTTAGCT

>Seq29_14_63294_0.0002

GCTATGTGAGACTAAAGTTAACT

>Seq30_14_63294_0.0002

GCTATGTGAGATTAAAGTTAA

>Seq31_13_63294_0.0002

GCTATGTGAGATTAAAGTCAACT

>Seq32_13_63294_0.0002

GCTATGTGAATTAAAGTTAACT

>Seq33_13_63294_0.0002

GCCATGTGAGATTAAAGTTAACT

>Seq34_12_63294_0.0002

ACTATGTGAGATTAAAGTTAACT

>Seq35_12_63294_0.0002

GCTATTTGAGATTAAAGTTAACT

>Seq36_12_63294_0.0002

GCTATGTGAGATTAAAGTTAACTC

>Seq37_11_63294_0.0002

GCTTGTGAGATTAAAGTTAACT

>Seq38_11_63294_0.0002

GCTATGTGAGATTAAAGCTAACT

>Seq39_11_63294_0.0002

GCTGTGTGAGATTAAAGTTAACT

>Seq40_11_63294_0.0002

GCTATGAGAGATTAAAGTTAACT

>Seq41_10_63294_0.0002

GCTATGTGAGATTGAAAGTTAACT

>Seq42_10_63294_0.0002

GCTAATGTGAGATTAAAGTTAACT

>Seq43_10_63294_0.0002

GCTATGTGAGATTAAAATTAACT

>Seq44_10_63294_0.0002

GCTATGTGAGATTAAGGTTAACT

>Seq45_10_63294_0.0002

GCTATGTGAGATTAAAGTAACT

>Seq46_9_63294_0.0001

GCTATGTTAGATTAAAGTTAACT

>Seq47_9_63294_0.0001

GCTATGTGAGGTTAAAGTTAACT

>Seq48_9_63294_0.0001

GCTATGTGAGATTAAAGTTAATT

>Seq49_9_63294_0.0001

GCTAGTGAGATTAAAGTTAACT

>Seq50_9_63294_0.0001

GCTATGTGATATTAAAGTTAACT

>Seq51_8_63294_0.0001

GCTATGTGAGATTAAAGTTAACTT

>Seq52_8_63294_0.0001

GCTATGTGAGATTTAAGTTAACT

>Seq53_8_63294_0.0001

GCTATGTAAGATTAAAGTTAACT

>Seq54_7_63294_0.0001

GCTATGTGAGATTAAAGTTAAAT

>Seq55_7_63294_0.0001

GCTATGTGAAGATTAAAGTTAACT

>Seq56_7_63294_0.0001

GCTATGTGAGTTTAAAGTTAACT

>Seq57_6_63294_0.0001

GCTATGTGAGATTAAAGTTTACT

>Seq58_6_63294_0.0001

GCTATGTGAGATTAAAGTTAACCT

>Seq59_6_63294_0.0001

GCTAAGTGAGATTAAAGTTAACT

>Seq60_6_63294_0.0001

GCTATGTGAGATTCCCTTTAAAGTTAACT

>Seq61_6_63294_0.0001

GCTATGTGAGATTAAAGTTAACG

>Seq62_5_63294_0.0001

GCTATGTGAGATTAAAGTAAACT

>Seq63_5_63294_0.0001

GCTATGTGAGATAAAGTTAACT

>Seq64_5_63294_0.0001

GCTTTGTGAGATTAAAGTTAACT

>Seq65_5_63294_0.0001

GCTATGTGTGATTAAAGTTAACT

>Seq66_5_63294_0.0001

GCTATGTGAGATAAAAGTTAACT

>Seq67_5_63294_0.0001

GCTATGTGAGAATAAAGTTAACT

>Seq68_5_63294_0.0001

GCTATGTGAGATTAAAGTTAAACT

>Seq69_4_63294_0.0001

GCTATGTGAGATTAAAGATAACT

>Seq70_4_63294_0.0001

GCTATGTGCGATTAAAGTTAACT

>Seq71_4_63294_0.0001

GCTATGTGAAATTAAAGTTAACT

>Seq72_4_63294_0.0001

GCTATGTGAGATGCAAAGTTAACT

>Seq73_3_63294_0.0000

GCTATGTGAGATTAAAGTTATCT

>Seq74_3_63294_0.0000

GCAATGTGAGATTAAAGTTAACT

>Seq75_3_63294_0.0000

GCTATGGTGAGATTAAAGTTAACT

>Seq76_3_63294_0.0000

GCTATGTGAGATTAATGTTAACT

>Seq77_3_63294_0.0000

GCTATGTGAGATTATAGTTAACT

>Seq78_3_63294_0.0000

GGTATGTGAGATTAAAGTTAACT

>Seq79_3_63294_0.0000

GCCTATGTGAGATTAAAGTTAACT

>Seq80_2_63294_0.0000

GCTCTGTGAGATTAAAGTTAACT

>Seq81_2_63294_0.0000

ACTATGTGAGATCAAAGTTAACT

>Seq82_2_63294_0.0000

GCTATGTGACATTAAAGTTAACT

>Seq83_2_63294_0.0000

GCTATGTGAGATTAAACTTAACT

>Seq84_2_63294_0.0000

GCTATGTGAGATTAAAGTTACCT

>Seq85_2_63294_0.0000

GCTATGTGAGATTAAAGTTTAACT

>Seq86_2_63294_0.0000

GCTATGGGAGATTAAAGTTAACT

>Seq87_2_63294_0.0000

GCTATGTGAGATTAAAGTTAT

>Seq88_2_63294_0.0000

GCTATGTGAGATTAACGTTAACT

>Seq89_2_63294_0.0000

GATATGTGAGATTAAAGTTAACT

>Seq90_2_63294_0.0000

GCTGTGAGATTAAAGTTAACT

>Seq91_2_63294_0.0000

GCTTATGTGAGATTAAAGTTAACT

>Seq92_2_63294_0.0000

CCTATGTGAGATTAAAGTTAACT

>Seq93_1_63294_0.0000

GCTATGTCAGATTAAAGTTAACT

>Seq94_1_63294_0.0000

GCTATGCGAGGTTAAAGTTAACT

>Seq95_1_63294_0.0000

GCTATGAGATTAAAGTTAACT

>Seq96_1_63294_0.0000

CTATGTGAGGTTAAAGTTAACT

>Seq97_1_63294_0.0000

GCTAGGTGAGATTAAAGTTAACT

>Seq98_1_63294_0.0000

GCTATGTGAGATTTAAAGTTAACC

>Seq99_1_63294_0.0000

GCTATGTGAGATTAAAGTACT

>Seq100_1_63294_0.0000

GCTATGTGAGATCAAAATTAACT

>Seq101_1_63294_0.0000

GCTATGTGAGATAAAGATAACT

>Seq102_1_63294_0.0000

GCTATGTTGAGATTAAAGTTAACT

>Seq103_1_63294_0.0000

GCTATGTGAGATTCAAAGTTAACT

>Seq104_1_63294_0.0000

GCTATGTGGATTAAAGTTAACT

>Seq105_1_63294_0.0000

GTATGTGAGATTAAGTTAACT

>Seq106_1_63294_0.0000

GCTATGTGAGTTTGAAAGTTAACT

>Seq107_1_63294_0.0000

GATGTGAGATTAAAGTTAACT

>Seq108_1_63294_0.0000

GCTATTGTGAGATTAAAGTTAACT

>Seq109_1_63294_0.0000

TTGCAGTGAGATTAAAGTTAACT

>Seq110_1_63294_0.0000

GCTATGTGAGGATTAAAGTTAACA

>Seq111_1_63294_0.0000

GCTATGTGAGGATCAAAGTTAACT

>Seq112_1_63294_0.0000

GCTATGTGAGATTAAAGCTATCT

>Seq113_1_63294_0.0000

GCTATCTGAGATTAAAGTTAACT

>Seq114_1_63294_0.0000

GCTATGTGAGAACGATGCAAAGTTAACT

>Seq115_1_63294_0.0000

GCTATGTGAGATTCCCTTTAAAGTTTACT

>Seq116_1_63294_0.0000

GCTATGTGAGAGTTAAAGTTAACT

>Seq117_1_63294_0.0000

GTTATGTGAGATTAAAGTTAAC

>Seq118_1_63294_0.0000

GCATGTGAGATTAAAGTTAAT

>Seq119_1_63294_0.0000

GCTATGTGAGAATTAAGGTTAACT

>Seq120_1_63294_0.0000

GCTAGTGAGGATTAAAGTTAACT

>Seq121_1_63294_0.0000

GCTATGTAGATTAAAGTTAACT

>Seq122_1_63294_0.0000

GGCTATGTGAGATTAAGTTAACT

>Seq123_1_63294_0.0000

GCATGTGAGGATTAAAGTTAACT

>Seq124_1_63294_0.0000

GCTATGTGAGATTACAGTTAACT

>Seq125_1_63294_0.0000

GCTATGTGAGACCAAAGTTAACT

>Seq126_1_63294_0.0000

GCTATGGAGATTAAAGTTAACT

>Seq127_1_63294_0.0000

GCATGTGAGATTAAAGTTAAC

>Seq128_1_63294_0.0000

GCTATTGAGATTAAAGTTAACT

>Seq129_1_63294_0.0000

GCGATGTGAGATTAAAGTTAACT
