## Supplementary Information files for "SARS-CoV-2 RdRp uses NDPs as a substrate and is able to incorporate NHC into RNA from diphosphate form molnupiravir": 13+23nt RNA NTP.docx

>Seq1_65960_67883_0.9717

GCTATGTGAGATTCCCTTTAAAGTTAACT

>Seq2_101_67883_0.0015

GCTATGTGAGATTCCCCTTTAAAGTTAACT

>Seq3_100_67883_0.0015

GCTATGTGAGATTCCCTTCAAAGTTAACT

>Seq4_96_67883_0.0014

GCTATGTGAGATTCCCTTAAAGTTAACT

>Seq5_81_67883_0.0012

GCTATGTGAGATTCCTTTAAAGTTAACT

>Seq6_65_67883_0.0010

GCTATGTGAGATTCCCTTTTAAAGTTAACT

>Seq7_63_67883_0.0009

GCTATGTGAGATTCCCTTTAAGTTAACT

>Seq8_60_67883_0.0009

GCTATGTGAGATTCCCTTTAAAGTTAAC

>Seq9_56_67883_0.0008

GCTATGTGAGATTTCCTTTAAAGTTAACT

>Seq10_54_67883_0.0008

CTATGTGAGATTCCCTTTAAAGTTAACT

>Seq11_40_67883_0.0006

GCTATGTGAGATTACCTTTAAAGTTAACT

>Seq12_40_67883_0.0006

GCTATGTGAGATTCCCTTTAAAGTTAACA

>Seq13_38_67883_0.0006

GCTATGTGAGATTCCCTTTAAAAGTTAACT

>Seq14_36_67883_0.0005

GCTATGTGAGATTACCCTTTAAAGTTAACT

>Seq15_35_67883_0.0005

GCTATGTGAGTTTCCCTTTAAAGTTAACT

>Seq16_32_67883_0.0005

GCTATGTGGGATTCCCTTTAAAGTTAACT

>Seq17_31_67883_0.0005

GCTATGTGAGATTTCCCTTTAAAGTTAACT

>Seq18_30_67883_0.0004

GCTATGTGAGATTCCCCTTAAAGTTAACT

>Seq19_29_67883_0.0004

GCTATGTGAGGATTCCCTTTAAAGTTAACT

>Seq20_26_67883_0.0004

GCTATGTGAGATTCCCTTTAAAGTTAACTC

>Seq21_24_67883_0.0004

GGCTATGTGAGATTCCCTTTAAAGTTAACT

>Seq22_24_67883_0.0004

GTTATGTGAGATTCCCTTTAAAGTTAACT

>Seq23_23_67883_0.0003

GCTATGTGAGATTCCCTTTAAAGTTAACC

>Seq24_23_67883_0.0003

GCTATGTGAGACTCCCTTTAAAGTTAACT

>Seq25_22_67883_0.0003

GCTATGTGAGATCCCCTTTAAAGTTAACT

>Seq26_20_67883_0.0003

GCTATGTGATATTCCCTTTAAAGTTAACT

>Seq27_20_67883_0.0003

GTATGTGAGATTCCCTTTAAAGTTAACT

>Seq28_20_67883_0.0003

GCTATGCGAGATTCCCTTTAAAGTTAACT

>Seq29_20_67883_0.0003

GCTATGTGAGATTCCCTTTAAAGTTAAT

>Seq30_19_67883_0.0003

GCTATGTGAGATTCACTTTAAAGTTAACT

>Seq31_18_67883_0.0003

GCTATGTGAGATTCCATTTAAAGTTAACT

>Seq32_17_67883_0.0003

GCTATGTGAGATTCCCTTTAAAGTTAGCT

>Seq33_17_67883_0.0003

GCTATGTGAGATTCCCTTTAAAGTTAA

>Seq34_16_67883_0.0002

GCTATGTGTGATTCCCTTTAAAGTTAACT

>Seq35_16_67883_0.0002

GCCATGTGAGATTCCCTTTAAAGTTAACT

>Seq36_16_67883_0.0002

GCTATATGAGATTCCCTTTAAAGTTAACT

>Seq37_16_67883_0.0002

GCTATGTGAGATTCCCTCTAAAGTTAACT

>Seq38_16_67883_0.0002

GCATGTGAGATTCCCTTTAAAGTTAACT

>Seq39_16_67883_0.0002

GCTACGTGAGATTCCCTTTAAAGTTAACT

>Seq40_16_67883_0.0002

GCTATGTGAGATTCCTTTTAAAGTTAACT

>Seq41_15_67883_0.0002

GCTATGTGAGAATTCCCTTTAAAGTTAACT

>Seq42_14_67883_0.0002

GCTATTTGAGATTCCCTTTAAAGTTAACT

>Seq43_13_67883_0.0002

TCTATGTGAGATTCCCTTTAAAGTTAACT

>Seq44_13_67883_0.0002

GCTATGTGAGATTCCCTTAAAAGTTAACT

>Seq45_12_67883_0.0002

GCTATGTGAGATTCCCTATAAAGTTAACT

>Seq46_12_67883_0.0002

GCTATGTAAGATTCCCTTTAAAGTTAACT

>Seq47_12_67883_0.0002

GCTTGTGAGATTCCCTTTAAAGTTAACT

>Seq48_12_67883_0.0002

GCTATGTGAGATTCCCTTTAAAGTCAACT

>Seq49_11_67883_0.0002

GCTATGTGAGATTCCCTTTAAAGTTACT

>Seq50_11_67883_0.0002

GCTATGTGAGGTTCCCTTTAAAGTTAACT

>Seq51_11_67883_0.0002

GCTATGTGAGATTCCCATTAAAGTTAACT

>Seq52_11_67883_0.0002

GCTATGTGAGATTCCCTTTGAAGTTAACT

>Seq53_11_67883_0.0002

GCTATGTGAGATTCCCTTTAAAATTAACT

>Seq54_10_67883_0.0001

GCTATGTTAGATTCCCTTTAAAGTTAACT

>Seq55_10_67883_0.0001

GCTATGTGAGATTCTCTTTAAAGTTAACT

>Seq56_10_67883_0.0001

GCTATGAGAGATTCCCTTTAAAGTTAACT

>Seq57_10_67883_0.0001

ACTATGTGAGATTCCCTTTAAAGTTAACT

>Seq58_9_67883_0.0001

GCTATGTGAGATTCCCTTTAGAGTTAACT

>Seq59_9_67883_0.0001

GCTATGTGAGATTCCCTTTAAGGTTAACT

>Seq60_9_67883_0.0001

GCTATGTGAGATTCCCTTTAAAGCTAACT

>Seq61_9_67883_0.0001

GCTAGTGAGATTCCCTTTAAAGTTAACT

>Seq62_9_67883_0.0001

GCTATGTGAGATTCCCTTTAAAGTTAACCT

>Seq63_9_67883_0.0001

GATATGTGAGATTCCCTTTAAAGTTAACT

>Seq64_8_67883_0.0001

GCTATGTGAGATTCCCTTTAAAGTTAATT

>Seq65_8_67883_0.0001

GCTATGTGAGATTCCCTTTAAAGTTAACTT

>Seq66_7_67883_0.0001

GCTGTGTGAGATTCCCTTTAAAGTTAACT

>Seq67_7_67883_0.0001

GCTATGTGAGATTAAAGTTAACT

>Seq68_7_67883_0.0001

GCTATGTGAGATACCCTTTAAAGTTAACT

>Seq69_7_67883_0.0001

GCTATGTGAGATTCCCTTTAAAGTAAACT

>Seq70_7_67883_0.0001

GCTATGTGAAATTCCCTTTAAAGTTAACT

>Seq71_7_67883_0.0001

GCTATGTGAGATTCCCTTTAAAGTTGACT

>Seq72_7_67883_0.0001

GCTATGTGAGATTAACCTTTAAAGTTAACT

>Seq73_6_67883_0.0001

GCTCTGTGAGATTCCCTTTAAAGTTAACT

>Seq74_6_67883_0.0001

GCTTTGTGAGATTCCCTTTAAAGTTAACT

>Seq75_6_67883_0.0001

GCCTATGTGAGATTCCCTTTAAAGTTAACT

>Seq76_6_67883_0.0001

GCTAAGTGAGATTCCCTTTAAAGTTAACT

>Seq77_6_67883_0.0001

GCTATGGGAGATTCCCTTTAAAGTTAACT

>Seq78_5_67883_0.0001

GCTATGTGAGATTCCCTTTAATGTTAACT

>Seq79_5_67883_0.0001

GCTATGTGGAGATTCCCTTTAAAGTTAACT

>Seq80_5_67883_0.0001

GCTATGTGAGATTCCCTTTAAAGTTTACT

>Seq81_5_67883_0.0001

GCTAATGTGAGATTCCCTTTAAAGTTAACT

>Seq82_4_67883_0.0001

GCTATGTGAGATTCCCTTTAAAGTTAAAT

>Seq83_4_67883_0.0001

GCTATGTGAGATTCCCTTTATAGTTAACT

>Seq84_4_67883_0.0001

GCTATGTGAGAATCCCTTTAAAGTTAACT

>Seq85_4_67883_0.0001

GCTATGTGAGATTAACCCTTTAAAGTTAACT

>Seq86_4_67883_0.0001

GCTATTGAGATTCCCTTTAAAGTTAACT

>Seq87_4_67883_0.0001

GCTATGTGAATTCCCTTTAAAGTTAACT

>Seq88_4_67883_0.0001

GCTATGTGAGATTGCCTTTAAAGTTAACT

>Seq89_3_67883_0.0000

GCTATGTGAGATTCCCTTTAAAGATAACT

>Seq90_3_67883_0.0000

GCTATCTGAGATTCCCTTTAAAGTTAACT

>Seq91_3_67883_0.0000

GCTATGTGAGATTCCCTTTAAAGTTAAACT

>Seq92_3_67883_0.0000

GATGTGAGATTCCCTTTAAAGTTAACT

>Seq93_3_67883_0.0000

GCTATGTGGATTCCCTTTAAAGTTAACT

>Seq94_3_67883_0.0000

GCTGTGAGATTCCCTTTAAAGTTAACT

>Seq95_3_67883_0.0000

GCTATGTGAGATTCCCTTTAAAGTTAACG

>Seq96_3_67883_0.0000

GCTATGTGACATTCCCTTTAAAGTTAACT

>Seq97_3_67883_0.0000

GCTATGTGAGAGTCCCTTTAAAGTTAACT

>Seq98_3_67883_0.0000

GCTATGTGAGATTCCCGTTAAAGTTAACT

>Seq99_3_67883_0.0000

GCTTATGTGAGATTCCCTTTAAAGTTAACT

>Seq100_2_67883_0.0000

GCTATGTGAGATTGCCCTTTAAAGTTAACT

>Seq101_2_67883_0.0000

GCTATGTGAGATTCCCTGAAAGTTAACT

>Seq102_2_67883_0.0000

GCTATGTGAGATTCCCTTTAAAGTTATCT

>Seq103_2_67883_0.0000

GCTATGTGAGATTCGCTTTAAAGTTAACT

>Seq104_2_67883_0.0000

CCTATGTGAGATTCCCTTTAAAGTTAACT

>Seq105_2_67883_0.0000

GCTATGTGCGATTCCCTTTAAAGTTAACT

>Seq106_2_67883_0.0000

GCTATGGTGAGATTCCCTTTAAAGTTAACT

>Seq107_2_67883_0.0000

GCTATGTGAGATTCTTTAAAGTTAACT

>Seq108_2_67883_0.0000

GCTATGTGAGATCCCTTTAAAGTTAACT

>Seq109_2_67883_0.0000

GCTATGTGAGATTACAAAAAGTTAACT

>Seq110_2_67883_0.0000

GCTATGTGAAGATTCCCTTTAAAGTTAACT

>Seq111_2_67883_0.0000

GGTGAGATTCCCTTTAAAGTTAACT

>Seq112_2_67883_0.0000

GCTATGTGAGATTCCCTTTAAAGTAACT

>Seq113_2_67883_0.0000

GCTATGTCAGATTCCCTTTAAAGTTAACT

>Seq114_2_67883_0.0000

GCTATGTGAGATTCCCTTAAGTTAACT

>Seq115_2_67883_0.0000

GCTATGTGAGATTCCCTCAAAGTTAACT

>Seq116_2_67883_0.0000

GCTATGTGAGATTCCCTTTAAAGGTAACT

>Seq117_2_67883_0.0000

GCTATGTGAGATTCCCTGTAAAGTTAACT

>Seq118_2_67883_0.0000

GCTATGGAGATTCCCTTTAAAGTTAACT

>Seq119_2_67883_0.0000

TATGTGAGATTCCCTTTAAAGTTAACT

>Seq120_1_67883_0.0000

GCTATGTGAGATTCCCTTTAAAGTTAT

>Seq121_1_67883_0.0000

GCTATGTGTAGATTCCCTTTAAAGTTAACT

>Seq122_1_67883_0.0000

GCTATGTGAGATTCCCTTTAAAGTGAATT

>Seq123_1_67883_0.0000

GCTATGTGAGATGAAAGTTAACT

>Seq124_1_67883_0.0000

GCTATGTGAGATTCCCCTTTAAAGTTAACG

>Seq125_1_67883_0.0000

GCTATGTGAGATTCCCTTTAACGTTAACT

>Seq126_1_67883_0.0000

GCTATGTGAGATTCCCTTTAAAGTTA

>Seq127_1_67883_0.0000

GCTATGTGAGATTCCATGCAAAGTTAACT

>Seq128_1_67883_0.0000

GCTATGTGAGATTCCCTTTAAGTTAGCT

>Seq129_1_67883_0.0000

GCTATGTGAGTTTTCCCTTTAAAGTTAACT

>Seq130_1_67883_0.0000

GGTATGTGAGATTCCCTTTAAAGTTAACT

>Seq131_1_67883_0.0000

GCTATGTGAGATTCCCTTAAAAGTTACT

>Seq132_1_67883_0.0000

GCTATGTGAGATTCCCTTTTAATTTAACT

>Seq133_1_67883_0.0000

GCTATGTGAGATTCCCTGCAAAGTTAACT

>Seq134_1_67883_0.0000

GCTATGTGATTCCCTTTAAAGTTAACT

>Seq135_1_67883_0.0000

GCTATGTGAGATTCCCTTTAAATTTAACT

>Seq136_1_67883_0.0000

ATGTGAGATTCCCTTTAAAGTTAACT

>Seq137_1_67883_0.0000

GCTATGTGAGATTCCCTTTAAAGTTTAACCT

>Seq138_1_67883_0.0000

GCTATGTTAGATTCCCTTTTAAAGTTAACT

>Seq139_1_67883_0.0000

GCTATGTGAGCTTCCCTTTAAAGTTAACT

>Seq140_1_67883_0.0000

GCTAGTGAGATTCCCTTTAAAGCTAACT

>Seq141_1_67883_0.0000

GCTATATGGGATTCCCTTTAAAGTTAACT

>Seq142_1_67883_0.0000

GTTATGTGAGATTCCTTTAAAGTTAACT

>Seq143_1_67883_0.0000

GTGTGAGATTCCCTTTAAAGTTAACT

>Seq144_1_67883_0.0000

GCTATGTGAGGTTCCCTTTAAAGTTAAC

>Seq145_1_67883_0.0000

GCTATGTGGGATTCCCCTTTAAAGTTAACT

>Seq146_1_67883_0.0000

GCTATGTGAGATTAAAAAAAAGTTAACT

>Seq147_1_67883_0.0000

GCTTTGTGTGATTCCCTTTAAAGTTAACT

>Seq148_1_67883_0.0000

GCTATGTGAGATCAAAGTTAACT

>Seq149_1_67883_0.0000

GCTATGTGAGATTTCCTTAAAGTTAACT

>Seq150_1_67883_0.0000

GCTATGTGAGATTCCCTTTTAAGTTAACT

>Seq151_1_67883_0.0000

GCTATGTGAGATTGACTTTAAAGTTAACT

>Seq152_1_67883_0.0000

GCTATGTGAGATTTTCCTTTAAAGTTAACT

>Seq153_1_67883_0.0000

GCTATGTGAGATTCCCTTTAAAGTTCACT

>Seq154_1_67883_0.0000

GCTATGTGAGTTTCCCTTTAAAAGTTAACT

>Seq155_1_67883_0.0000

GCTATGTGAGATTTCACTTTAAAGTTAACT

>Seq156_1_67883_0.0000

GCTTGTGAGACTCCCTTTAAAGTTAACT

>Seq157_1_67883_0.0000

GCTATTGTGAGATTCCCTTTAAAGTTAACT

>Seq158_1_67883_0.0000

GCTATGTGAGATTCCCCTTTAAAGTTAA

>Seq159_1_67883_0.0000

GCTATGTGAGATACCGACTTTAAAGTTAACT

>Seq160_1_67883_0.0000

CTATGTGAGATTCCTTTAAAGTTAACT

>Seq161_1_67883_0.0000

GCTATGTGAGATTCCCTTTAAAGTTAACTCCGCTCGAGCGGTGAATTGGTATCGACCATTTGTCGAGACTTCCCTTTAAAGTTAACT

>Seq162_1_67883_0.0000

GCTATGTGAGATTCCCTTTTAAAGCTAACT

>Seq163_1_67883_0.0000

GCTATGTGAGGATTCCCCTTTAAAGTTAACT

>Seq164_1_67883_0.0000

GCATGTGAGATTCCCTTTAAAGTTACT

>Seq165_1_67883_0.0000

GCATGTGAGATTCCCTTTAAAGATAACT

>Seq166_1_67883_0.0000

GCTATGTGAGATGCCCTTTAAAGTTAACT

>Seq167_1_67883_0.0000

GCTATGTGAGATTTTTAAAGTTAACT

>Seq168_1_67883_0.0000

GCTATGTGAGATTCCCTTTACAGTTAACT

>Seq169_1_67883_0.0000

GCGATGTGAGATTCCCTTTAAAGTTAACT

>Seq170_1_67883_0.0000

GCTATGTGAGATTCCCTAAAGTTAACT

>Seq171_1_67883_0.0000

GCAATGTGAGATTCCCTTTAAAGTTAACT

>Seq172_1_67883_0.0000

GCTATGTGAGATTCCCTTGAAAGTTAACT

>Seq173_1_67883_0.0000

GCTATGTGAGATTCCCTTCAAAGTTAAC

>Seq174_1_67883_0.0000

GCTATGTGAGATTCCCCTTTAAAGTTAAACT

>Seq175_1_67883_0.0000

GCTATGTGAGATTCCCATAAAGTTAACT

>Seq176_1_67883_0.0000

GCTATCGAGATTCCCTTTAAAGTTAACT

>Seq177_1_67883_0.0000

GCTATGTGAGATTCCCCTGAAAGTTAACT

>Seq178_1_67883_0.0000

GCTATGTGAGATTCTAAAGTTAACT

>Seq179_1_67883_0.0000

GCTATGTGAGATTCCTTAAAGTTAACT

>Seq180_1_67883_0.0000

GCTATGTGAGGATTCCTTTAAAGTTAACT

>Seq181_1_67883_0.0000

GCTATGTGAGATTCCCTTTAAAGGTTAACT

>Seq182_1_67883_0.0000

GCATGTGAGATTCCCTTTAGAGTTAACT
