## Supplementary Information files for "SARS-CoV-2 RdRp uses NDPs as a substrate and is able to incorporate NHC into RNA from diphosphate form molnupiravir": 13+29nt RNA NTP.docx

No mutations: TCCCTTT	67883-778=67105 98.85%

Mutations: 778

1: TCCCCTTT	128

2: TCCCTTC	111

3: TTCCTTT	157

4: TACCTTT	40

5: TACCCTTT	43

6: TCCCCTT	60

7: TCACTTT	20

8: TCCATTT	18

9: TCCCTCT	16

10: TCCCTAT	12

11: TCCCATT	11

12: TCTCTTT	10

13: TAACCTTT	7

14: TAACCCTTT	4

15: TGCCTTT	4

16: TCCCGTT	3

17: TGCCCTTT	3

18: TCCCTG	5

19: TCGCTTT	2

20: TCTTT	2

21: TCCCTC	2

22: TCCATGC	1

23: TCCCTTA	112

24: TAAAAA	1

25: TTCCTT	2

26: ACCGACTTT	1

27: TCCCTTG	1

28: TCCCAT	1

29: TCCCCTG	1
