## Supplementary Information files for "SARS-CoV-2 RdRp uses NDPs as a substrate and is able to incorporate NHC into RNA from diphosphate form molnupiravir": 50 μM CTP 50 μM MDP.docx

No mutation: TCCCTTT	61673-619=61054	99.00%

Mutations: 619

1: TCCCCTTT	119

2: TCCCTTC	85

3: TACCCTTT	39

4: TCCCCTT	43

5: TCCTTTT	22

6: TACCTTT	23

7: TCCCTCT	15

8: TCTCTTT	11

9: TCCATTT	28

10: TCCCTTA	69

11: TCACTTT	18

12: TCCCTGC	1

13: TCGCTTT	1

14: TCCCTAT	9

15: TCCCATT	10

16: TAACCCTTT	5

17: TCTTT	4

18: TGCCTTT	3

19: TCCTTT	104

20: TAAAAA	2

21: TCCCTTG	3

22: TCCAAA	2

23: TCCCTG	3

>Seq1_59914_61673_0.9715

GCTATGTGAGATTCCCTTTAAAGTTAACT

>Seq2_96_61673_0.0016

GCTATGTGAGATTCCCCTTTAAAGTTAACT

>Seq3_84_61673_0.0014

GCTATGTGAGATTCCCTTCAAAGTTAACT

>Seq4_78_61673_0.0013

GCTATGTGAGATTCCTTTAAAGTTAACT

>Seq5_70_61673_0.0011

GCTATGTGAGATTCCCTTTAAGTTAACT

>Seq6_67_61673_0.0011

GCTATGTGAGATTCCCTTTTAAAGTTAACT

>Seq7_61_61673_0.0010

GCTATGTGAGATTCCCTTTAAAGTTAAC

>Seq8_58_61673_0.0009

GCTATGTGAGATTCCCTTAAAGTTAACT

>Seq9_49_61673_0.0008

CTATGTGAGATTCCCTTTAAAGTTAACT

>Seq10_46_61673_0.0007

GCTATGTGAGATTTCCTTTAAAGTTAACT

>Seq11_42_61673_0.0007

GCTATGTGAGATTCCCCTTAAAGTTAACT

>Seq12_41_61673_0.0007

GCTATGTGAGATTTCCCTTTAAAGTTAACT

>Seq13_32_61673_0.0005

GCTATGTGAGATTACCCTTTAAAGTTAACT

>Seq14_29_61673_0.0005

GCTATGTGAGTTTCCCTTTAAAGTTAACT

>Seq15_29_61673_0.0005

GCTATGTGAGGATTCCCTTTAAAGTTAACT

>Seq16_27_61673_0.0004

GCTATGTGAGATTCCCTTTAAAGTTAACA

>Seq17_27_61673_0.0004

GCTATGTGAGATTCCATTTAAAGTTAACT

>Seq18_27_61673_0.0004

GCTATGTGGGATTCCCTTTAAAGTTAACT

>Seq19_24_61673_0.0004

GCTATGTGAGATTCCCTTTAAAAGTTAACT

>Seq20_23_61673_0.0004

GCTATGCGAGATTCCCTTTAAAGTTAACT

>Seq21_22_61673_0.0004

GCTATGTGAGATTACCTTTAAAGTTAACT

>Seq22_22_61673_0.0004

GCTATGTGAGATTCCTTTTAAAGTTAACT

>Seq23_21_61673_0.0003

GCTATGTGAGATCCCCTTTAAAGTTAACT

>Seq24_20_61673_0.0003

GGCTATGTGAGATTCCCTTTAAAGTTAACT

>Seq25_20_61673_0.0003

GCTACGTGAGATTCCCTTTAAAGTTAACT

>Seq26_19_61673_0.0003

GCTATATGAGATTCCCTTTAAAGTTAACT

>Seq27_19_61673_0.0003

GCTATGTGAGATTCCCTTTAAAGTCAACT

>Seq28_18_61673_0.0003

GTTATGTGAGATTCCCTTTAAAGTTAACT

>Seq29_18_61673_0.0003

GCTATGTGAGATTCACTTTAAAGTTAACT

>Seq30_17_61673_0.0003

GCTATGTGAGACTCCCTTTAAAGTTAACT

>Seq31_16_61673_0.0003

GCTATGTGAGATTCCCTTTAAAGTTAACC

>Seq32_16_61673_0.0003

GCTATGTGAGGTTCCCTTTAAAGTTAACT

>Seq33_15_61673_0.0002

GTATGTGAGATTCCCTTTAAAGTTAACT

>Seq34_15_61673_0.0002

GCTATGTGAGATTCCCTTTAAAGTTAACTC

>Seq35_15_61673_0.0002

GCTATGTGTGATTCCCTTTAAAGTTAACT

>Seq36_15_61673_0.0002

GCTATGTGAGATTCCCTCTAAAGTTAACT

>Seq37_14_61673_0.0002

GCTATGTTAGATTCCCTTTAAAGTTAACT

>Seq38_14_61673_0.0002

GCATGTGAGATTCCCTTTAAAGTTAACT

>Seq39_14_61673_0.0002

ACTATGTGAGATTCCCTTTAAAGTTAACT

>Seq40_13_61673_0.0002

GCTATGTAAGATTCCCTTTAAAGTTAACT

>Seq41_13_61673_0.0002

TCTATGTGAGATTCCCTTTAAAGTTAACT

>Seq42_13_61673_0.0002

GCTATGTGAGATTCCCTTTAAGGTTAACT

>Seq43_13_61673_0.0002

GCCATGTGAGATTCCCTTTAAAGTTAACT

>Seq44_13_61673_0.0002

GCTATGTGAGATTCCCTTTAAAGTTAAT

>Seq45_13_61673_0.0002

GCTATGTGAGATTCCCTTTAAAGTTGACT

>Seq46_12_61673_0.0002

GCTATGTGAGATTCCCTTTAAAGTTAGCT

>Seq47_12_61673_0.0002

GCTTGTGAGATTCCCTTTAAAGTTAACT

>Seq48_12_61673_0.0002

GCTATTTGAGATTCCCTTTAAAGTTAACT

>Seq49_11_61673_0.0002

GCTAGTGAGATTCCCTTTAAAGTTAACT

>Seq50_11_61673_0.0002

GCTATGTGAGATTCTCTTTAAAGTTAACT

>Seq51_11_61673_0.0002

GCTATGTGAGATTCCCTTTAAAGTTAACCT

>Seq52_11_61673_0.0002

GCTATGTGAGATTCCCTTTAAAGTTAA

>Seq53_11_61673_0.0002

GCTATGTGAGATTCCCTTAAAAGTTAACT

>Seq54_11_61673_0.0002

GCTATGTGAGATTCCCTTTAAAATTAACT

>Seq55_10_61673_0.0002

GCTATGTGACATTCCCTTTAAAGTTAACT

>Seq56_10_61673_0.0002

GCTATGTGAGATTCCCTTTAAAGTTTACT

>Seq57_9_61673_0.0001

GCTATGTGATATTCCCTTTAAAGTTAACT

>Seq58_9_61673_0.0001

GCTATGTGAGAATCCCTTTAAAGTTAACT

>Seq59_9_61673_0.0001

GCTATGTGAGATTCCCTTTAAAGCTAACT

>Seq60_9_61673_0.0001

GCTATGTGAGATTCCCATTAAAGTTAACT

>Seq61_9_61673_0.0001

GCTATGTGAGATTCCCTTTTAAGTTAACT

>Seq62_8_61673_0.0001

GCTATGTGAGATTCCCTATAAAGTTAACT

>Seq63_8_61673_0.0001

GCTATGTGAGATTCCCTTTGAAGTTAACT

>Seq64_8_61673_0.0001

GCTATGTGAGATTCCCTTTAAAGTTAATT

>Seq65_8_61673_0.0001

GCTATGTGAAATTCCCTTTAAAGTTAACT

>Seq66_7_61673_0.0001

GCTATGTGAGATTCCCTTTAGAGTTAACT

>Seq67_7_61673_0.0001

GCTATGTGAGAATTCCCTTTAAAGTTAACT

>Seq68_7_61673_0.0001

GCTATGTGAGATTCCCTTTAAAGTTACT

>Seq69_7_61673_0.0001

GCTATGTGCGATTCCCTTTAAAGTTAACT

>Seq70_7_61673_0.0001

GCTGTGAGATTCCCTTTAAAGTTAACT

>Seq71_7_61673_0.0001

GCTATGAGAGATTCCCTTTAAAGTTAACT

>Seq72_6_61673_0.0001

GCTATGTGAGATTCCCTTTATAGTTAACT

>Seq73_6_61673_0.0001

GCTATGTGAGATTCCCTTTAATGTTAACT

>Seq74_6_61673_0.0001

GCTATGTGAGATACCCTTTAAAGTTAACT

>Seq75_6_61673_0.0001

GCTATGTGAGATTCCCTTTAAAGTTAACG

>Seq76_6_61673_0.0001

GCTAAGTGAGATTCCCTTTAAAGTTAACT

>Seq77_5_61673_0.0001

GCTATGTGAGATTAACCCTTTAAAGTTAACT

>Seq78_4_61673_0.0001

GCTGTGTGAGATTCCCTTTAAAGTTAACT

>Seq79_4_61673_0.0001

GCTATGTGAGATTCCCTTTAAAGTAACT

>Seq80_4_61673_0.0001

GCTATGTGAGATTCCCGTTAAAGTTAACT

>Seq81_3_61673_0.0000

GCTATGTGAGATTCCCTTTAAAGTTATCT

>Seq82_3_61673_0.0000

GGTATGTGAGATTCCCTTTAAAGTTAACT

>Seq83_3_61673_0.0000

GCTATGTGAGATTCCCTTTAAAGTTAAACT

>Seq84_3_61673_0.0000

GCTATGTGGATTCCCTTTAAAGTTAACT

>Seq85_3_61673_0.0000

GCTATGTGAGATTAAAGTTAACT

>Seq86_3_61673_0.0000

GCTATGTGAGATTCCCTTTCAAGTTAACT

>Seq87_3_61673_0.0000

GCTATGTGAGATCCCTTTAAAGTTAACT

>Seq88_3_61673_0.0000

GCTATGTGAAGATTCCCTTTAAAGTTAACT

>Seq89_3_61673_0.0000

GCCTATGTGAGATTCCCTTTAAAGTTAACT

>Seq90_3_61673_0.0000

GATATGTGAGATTCCCTTTAAAGTTAACT

>Seq91_3_61673_0.0000

GCGATGTGAGATTCCCTTTAAAGTTAACT

>Seq92_3_61673_0.0000

GCTATGGGAGATTCCCTTTAAAGTTAACT

>Seq93_3_61673_0.0000

GCTATGTGAGATTCCCTTGAAAGTTAACT

>Seq94_3_61673_0.0000

GCTAATGTGAGATTCCCTTTAAAGTTAACT

>Seq95_3_61673_0.0000

GCTATGTGAGATTGCCTTTAAAGTTAACT

>Seq96_2_61673_0.0000

GCTATGTGAGATTCCCTTTAAAGTTAT

>Seq97_2_61673_0.0000

GCTATGTGAGATTCCCTTTAAAGTTAAAT

>Seq98_2_61673_0.0000

GCTATGTTGAGATTCCCTTTAAAGTTAACT

>Seq99_2_61673_0.0000

GCTATGTGAGATTCCCTTTAAAGATAACT

>Seq100_2_61673_0.0000

GCTATGTGAGATTCCCTTTAAATTTAACT

>Seq101_2_61673_0.0000

ATGTGAGATTCCCTTTAAAGTTAACT

>Seq102_2_61673_0.0000

GCTATGTGAGATTCAAAGTTAACT

>Seq103_2_61673_0.0000

GCTATGTGAGATTCCCTCCAAAGTTAACT

>Seq104_2_61673_0.0000

GCTATGTGGAGATTCCCTTTAAAGTTAACT

>Seq105_2_61673_0.0000

GCTATGGTGAGATTCCCTTTAAAGTTAACT

>Seq106_2_61673_0.0000

GCTATGTGAGATTCTTTAAAGTTAACT

>Seq107_2_61673_0.0000

GCTTTGTGAGATTCCCTTTAAAGTTAACT

>Seq108_2_61673_0.0000

GCTATGTCAGATTCCCTTTAAAGTTAACT

>Seq109_2_61673_0.0000

GCTATGTGAGATTCCCTCAAAGTTAACT

>Seq110_2_61673_0.0000

GCTATGTGAGATTCTTAAAGTTAACT

>Seq111_2_61673_0.0000

GCTATGTGAGATTCCCTTTACAGTTAACT

>Seq112_2_61673_0.0000

GCTATGTGAGATTCCCTTTAAAGTTACCT

>Seq113_2_61673_0.0000

GCTATGGAGATTCCCTTTAAAGTTAACT

>Seq114_2_61673_0.0000

GCAATGTGAGATTCCCTTTAAAGTTAACT

>Seq115_2_61673_0.0000

GCTATGTAGATTCCCTTTAAAGTTAACT

>Seq116_2_61673_0.0000

GCTATGTGAGATTCCCTTTAAAGTTAACTT

>Seq117_2_61673_0.0000

GCTATGTGAGATTCCCTTTAAAGTTTAACT

>Seq118_2_61673_0.0000

GCTTATGTGAGATTCCCTTTAAAGTTAACT

>Seq119_1_61673_0.0000

GCTATGTGAGATTCCCTTTAAAGTTACAT

>Seq120_1_61673_0.0000

GCTATGTGAGATTCCCTATTAAAGTTAACT

>Seq121_1_61673_0.0000

GCTATGTGAGATTCCCTTTTAAATTAACT

>Seq122_1_61673_0.0000

GCTATGTGAGATTCCCTGAAAGTTAACT

>Seq123_1_61673_0.0000

GCTATGTGAGATTCGCTTTAAAGTTAACT

>Seq124_1_61673_0.0000

GTGTGAGATCCCCTTTAAAGTTAACT

>Seq125_1_61673_0.0000

GCTATGAGATTCCCTTTAAAGTTAACT

>Seq126_1_61673_0.0000

GCTATGTGGGATTCCTTTAAAGTTAACT

>Seq127_1_61673_0.0000

GCTATGTGAGTTCCCTTTAAAGTTAACT

>Seq128_1_61673_0.0000

GCTATGTGAGATTCCCTGGAAAGTTAACT

>Seq129_1_61673_0.0000

GCTCTATGAGATTCCCTTTAAAGTTAACT

>Seq130_1_61673_0.0000

GCTATCTGAGATTCCCTTTAAAGTTAACT

>Seq131_1_61673_0.0000

GGCTATGTGAGTTTCCCTTTAAAGTTAACT

>Seq132_1_61673_0.0000

GCTATGTGAGATTCCTTTAAAGTTAAT

>Seq133_1_61673_0.0000

GCTATGTGAGATTCCCTTTAAAGTTAAGT

>Seq134_1_61673_0.0000

GCTATGTGAGATTCCCTGCAAAGTTAACT

>Seq135_1_61673_0.0000

GCTATGTAAAGATTCCCTTTAAAGTTAACT

>Seq136_1_61673_0.0000

GCTATGTGAGGATTCCCTTTAAAGTTAAC

>Seq137_1_61673_0.0000

GCTCTTTTAGATTCCCTTTAAAGTTAACT

>Seq138_1_61673_0.0000

TGTGAGATTCCCTTTAAAGTTAACT

>Seq139_1_61673_0.0000

GTGTGAGATTCCCTTTAAAGTTAACT

>Seq140_1_61673_0.0000

GCTAGGTGAGATTCCCTTTAAAGTTAACT

>Seq141_1_61673_0.0000

GATGTGAGATTCCCTTTAAAGTTAACT

>Seq142_1_61673_0.0000

GCTATGTGAGATTACCCTTTAAAGTTAACA

>Seq143_1_61673_0.0000

CTATGTGAGATTCCCTTTAAAGTTAACCT

>Seq144_1_61673_0.0000

GCTATGTGAGATTCCCTTTAAGTTAACCT

>Seq145_1_61673_0.0000

GCTATGTGAGATTACCCCTTTAAAGTTAACT

>Seq146_1_61673_0.0000

GCTCTGTGAGATTCCCTTTAAAGTTAACT

>Seq147_1_61673_0.0000

GCTATGTGAGATTCCCTTTAAAGTTAACTCT

>Seq148_1_61673_0.0000

GCTATGTGAGATTAAAAAAAAAGTTAACT

>Seq149_1_61673_0.0000

GCTATGTGAGATTCCCTAAGTTAACT

>Seq150_1_61673_0.0000

GCTATGTGAGATTATCTTTAAAGTTAACT

>Seq151_1_61673_0.0000

GGTATGAGAGATTCCCTTTAAAGTTAACT

>Seq152_1_61673_0.0000

GCTATGTGAGATTACCATTAAAGTTAACT

>Seq153_1_61673_0.0000

GCTATTGTGAGATTCCCTTTAAAGTTAACT

>Seq154_1_61673_0.0000

GCGTGAGATTCCCTTTAAAGTTAACT

>Seq155_1_61673_0.0000

GCTATGTGAGATTTTAAAGTTAACT

>Seq156_1_61673_0.0000

GCTATGTGAGATTCCCATTTAAAGTTAACT

>Seq157_1_61673_0.0000

GCTATGTGAGATTCCCTTTAAAGTAAACT

>Seq158_1_61673_0.0000

GCTATGTGAGATTCCTCTTTAAAGTTAACT

>Seq159_1_61673_0.0000

GCTCTGTGAGATTCCCTTTAAAGTTACCT

>Seq160_1_61673_0.0000

GCTATGTGAGATTCCATTTAAAGTAACT

>Seq161_1_61673_0.0000

GCTATGTGAGATTTTTAAAGTTAACT

>Seq162_1_61673_0.0000

GTTATGTGAGATTCCCCTTTAAAGTTAACT

>Seq163_1_61673_0.0000

GCTATGTGAGATTCCCTTTAAAGGTAACT

>Seq164_1_61673_0.0000

GCTATGTGAGATTATCCTTTAAAGTTAACT

>Seq165_1_61673_0.0000

GCTATGTGAGATTACCTTAAAGTTAACT

>Seq166_1_61673_0.0000

GCTATGTGAGATTCCCTTCAAAGTTAAT

>Seq167_1_61673_0.0000

GCTATGTGGGATTAACCTTTAAAGTTAACT

>Seq168_1_61673_0.0000

GCTATTGAGATTCCCTTTAAAGTTAACT

>Seq169_1_61673_0.0000

GCTATGTGAATTCCCTTTAAAGTTAACT

>Seq170_1_61673_0.0000

GCTATGTGAGATTCCCCTTAAAAGTTAACT

>Seq171_1_61673_0.0000

GCTATGTGAGATTTCCTTTAAAGTTAA

>Seq172_1_61673_0.0000

GCTATGTGAGATTCCCTAAAGTTAACT

>Seq173_1_61673_0.0000

TCTATGTGAGATTACCTTTAAAGTTAACT

>Seq174_1_61673_0.0000

GCTATGTGAGATTAACCTTTAAAGTTAACT

>Seq175_1_61673_0.0000

GCTATGTGAGATTAAAAAAAGTTAACT

>Seq176_1_61673_0.0000

GCTATGTGAGATTCCCCAAAGTTAACT

>Seq177_1_61673_0.0000

GCTATGTGAGATTCCTTAAAGTTAACT

>Seq178_1_61673_0.0000

TATGTGAGATTCCCTTTAAAGTTAACT

>Seq179_1_61673_0.0000

GCTATGTGAGATTCCTCTAAAGTTAACT

>Seq180_1_61673_0.0000

CTATGTGAGATTCCCTTTTAAAGTTAACT
