## Supplementary Information files for "SARS-CoV-2 RdRp uses NDPs as a substrate and is able to incorporate NHC into RNA from diphosphate form molnupiravir": 50 μM CTP 50 μM MTP.docx

No mutation: TCCCTTT	64316-690=63626 98.93%

Mutation: 690

1: TCCCCTTT	150

2: TCCCTTC	102

3: TACCCTTT	40

4: TCCCCTT	32

5: TCCTTT T	26

6: TACCTTT	22

7: TCCCTCT	17

8: TCTCTTT	17

9: TCCATTT	16

10: TCCCTTA	88

11: TCACTTT	12

12: TCCCTGC	7

13: TCGCTTT	4

14: TCCCTAT	4

15: TCCCATT	4

16: TAACCCTTT	5

17: TCTTT	4

18: TGCCTTT	4

19: TCCCCTC	2

20: GCCCTTT	3

21: TCCCTTG	2

22: TCCAAA	2

23: TCCCTG	2

24: TCTCTTC	1

25: TCCCTCAT	1

26: TCCTTT	121

27: TAAAAA	1

28: TTCCTTC	1

29: TCCGTTT	1

>Seq1_62551_64316_0.9726

GCTATGTGAGATTCCCTTTAAAGTTAACT

>Seq2_117_64316_0.0018

GCTATGTGAGATTCCCCTTTAAAGTTAACT

>Seq3_102_64316_0.0016

GCTATGTGAGATTCCCTTCAAAGTTAACT

>Seq4_75_64316_0.0012

GCTATGTGAGATTCCCTTAAAGTTAACT

>Seq5_74_64316_0.0012

GCTATGTGAGATTCCTTTAAAGTTAACT

>Seq6_63_64316_0.0010

GCTATGTGAGATTCCCTTTTAAAGTTAACT

>Seq7_57_64316_0.0009

CTATGTGAGATTCCCTTTAAAGTTAACT

>Seq8_55_64316_0.0009

GCTATGTGAGATTCCCTTTAAGTTAACT

>Seq9_45_64316_0.0007

GCTATGTGAGATTCCCTTTAAAAGTTAACT

>Seq10_43_64316_0.0007

GCTATGTGAGATTTCCCTTTAAAGTTAACT

>Seq11_42_64316_0.0007

GCTATGTGAGATTTCCTTTAAAGTTAACT

>Seq12_42_64316_0.0007

GCTATGTGAGATTCCCTTTAAAGTTAAC

>Seq13_36_64316_0.0006

GCTATGTGAGATTACCCTTTAAAGTTAACT

>Seq14_36_64316_0.0006

GCTATGTGGGATTCCCTTTAAAGTTAACT

>Seq15_35_64316_0.0005

GCTATGTGAGTTTCCCTTTAAAGTTAACT

>Seq16_33_64316_0.0005

GCTATGTGAGGATTCCCTTTAAAGTTAACT

>Seq17_30_64316_0.0005

GCTATGTGAGATTCCCCTTAAAGTTAACT

>Seq18_28_64316_0.0004

GCTATGTGAGATTCCCTTTAAAGTTAACA

>Seq19_25_64316_0.0004

GTTATGTGAGATTCCCTTTAAAGTTAACT

>Seq20_24_64316_0.0004

GCTATGTGAGATTCCCTTTAAAGTTAACC

>Seq21_24_64316_0.0004

GCTATGTGAGATTCCTTTTAAAGTTAACT

>Seq22_22_64316_0.0003

GCTATGTGAGATTACCTTTAAAGTTAACT

>Seq23_19_64316_0.0003

GCCATGTGAGATTCCCTTTAAAGTTAACT

>Seq24_18_64316_0.0003

GCTATGTGAGATTCCCTTTAAAGTTAACTC

>Seq25_18_64316_0.0003

GCTATGTGAGATCCCCTTTAAAGTTAACT

>Seq26_17_64316_0.0003

GGCTATGTGAGATTCCCTTTAAAGTTAACT

>Seq27_17_64316_0.0003

GCTATGTGAGATTCCCTCTAAAGTTAACT

>Seq28_17_64316_0.0003

GCTATGTGAGACTCCCTTTAAAGTTAACT

>Seq29_16_64316_0.0002

GCTATGTGATATTCCCTTTAAAGTTAACT

>Seq30_16_64316_0.0002

GCTATTTGAGATTCCCTTTAAAGTTAACT

>Seq31_16_64316_0.0002

GCTATGTGAGATTCTCTTTAAAGTTAACT

>Seq32_15_64316_0.0002

GCTATGTGTGATTCCCTTTAAAGTTAACT

>Seq33_15_64316_0.0002

GCTATATGAGATTCCCTTTAAAGTTAACT

>Seq34_15_64316_0.0002

GCTATGCGAGATTCCCTTTAAAGTTAACT

>Seq35_14_64316_0.0002

GTATGTGAGATTCCCTTTAAAGTTAACT

>Seq36_14_64316_0.0002

GCTATGTGAGATTCCCTTTAAGGTTAACT

>Seq37_14_64316_0.0002

GCTATGTGAGATTCCATTTAAAGTTAACT

>Seq38_14_64316_0.0002

GCTATGTGAGATTCCCTTTAAAGTTAAT

>Seq39_13_64316_0.0002

GCTATGTGAGATTCCCTTTAAAGCTAACT

>Seq40_13_64316_0.0002

GCTATGTGAGATTCCCTTTAAAGTTAA

>Seq41_13_64316_0.0002

GCTATGTGAGATTCCCTTAAAAGTTAACT

>Seq42_13_64316_0.0002

GCATGTGAGATTCCCTTTAAAGTTAACT

>Seq43_13_64316_0.0002

GCTACGTGAGATTCCCTTTAAAGTTAACT

>Seq44_12_64316_0.0002

GCTATGTAAGATTCCCTTTAAAGTTAACT

>Seq45_12_64316_0.0002

GCTATGTGAGATTCACTTTAAAGTTAACT

>Seq46_12_64316_0.0002

GCTATGTGAGATTCCCTTTAAAGTTAATT

>Seq47_12_64316_0.0002

GCTATGTGAGATTCCCTTTAAAGTTAACCT

>Seq48_11_64316_0.0002

GCTATGTGAGAATTCCCTTTAAAGTTAACT

>Seq49_11_64316_0.0002

GCTGTGTGAGATTCCCTTTAAAGTTAACT

>Seq50_11_64316_0.0002

GCTATGTGAGGTTCCCTTTAAAGTTAACT

>Seq51_11_64316_0.0002

GCTATGTGAAATTCCCTTTAAAGTTAACT

>Seq52_10_64316_0.0002

GCTATGTTAGATTCCCTTTAAAGTTAACT

>Seq53_9_64316_0.0001

GCTTGTGAGATTCCCTTTAAAGTTAACT

>Seq54_9_64316_0.0001

GCTATGTGAGATTCCCTTTTAAGTTAACT

>Seq55_8_64316_0.0001

GCTATGTGAGATTCCCTTTAGAGTTAACT

>Seq56_8_64316_0.0001

GCTATGTGAGATTCCCTTTAATGTTAACT

>Seq57_8_64316_0.0001

GCTATGTGAGATTCCCTTTAAAGTTAGCT

>Seq58_8_64316_0.0001

GCTATGTGAGATTCCCTTTAAAGTTACT

>Seq59_8_64316_0.0001

TCTATGTGAGATTCCCTTTAAAGTTAACT

>Seq60_8_64316_0.0001

GCTAGTGAGATTCCCTTTAAAGTTAACT

>Seq61_8_64316_0.0001

GCTATGTGAGATTCCCTTTAAAGTAAACT

>Seq62_7_64316_0.0001

GCTATGTGAGAATCCCTTTAAAGTTAACT

>Seq63_7_64316_0.0001

GCTATGTGAGATTCCCTGCAAAGTTAACT

>Seq64_7_64316_0.0001

GCTATGTGAGATTCCCTTTAAAGTCAACT

>Seq65_7_64316_0.0001

ACTATGTGAGATTCCCTTTAAAGTTAACT

>Seq66_6_64316_0.0001

GCTTTGTGAGATTCCCTTTAAAGTTAACT

>Seq67_6_64316_0.0001

GCTATGTGACATTCCCTTTAAAGTTAACT

>Seq68_6_64316_0.0001

GCTATGGGAGATTCCCTTTAAAGTTAACT

>Seq69_6_64316_0.0001

GCTATGTGAATTCCCTTTAAAGTTAACT

>Seq70_6_64316_0.0001

GCTATGTGAGATTCCCTTTAAAGTTGACT

>Seq71_5_64316_0.0001

GCTATCTGAGATTCCCTTTAAAGTTAACT

>Seq72_5_64316_0.0001

GCTCTGTGAGATTCCCTTTAAAGTTAACT

>Seq73_5_64316_0.0001

GCTATGTGAGATTCCCTTTAAAGTTTACT

>Seq74_5_64316_0.0001

GCTATGAGAGATTCCCTTTAAAGTTAACT

>Seq75_5_64316_0.0001

GCTATGTGAGATTCCCTTTAAAGTTAACTT

>Seq76_5_64316_0.0001

GCTATGTGAGATTCCCTTTAAAATTAACT

>Seq77_4_64316_0.0001

GCTATGTGAGATTCCCTTTAAAGTTAAAT

>Seq78_4_64316_0.0001

GCTATGTGAGATTCCCTTTATAGTTAACT

>Seq79_4_64316_0.0001

GCTATGTGAGATTCGCTTTAAAGTTAACT

>Seq80_4_64316_0.0001

GCTATGTGAGATTCCCTATAAAGTTAACT

>Seq81_4_64316_0.0001

GCTATGTGAGATTCCCTTTAAAGATAACT

>Seq82_4_64316_0.0001

GCTATGTGAGATTCCCATTAAAGTTAACT

>Seq83_4_64316_0.0001

GCTATGTGAGATACCCTTTAAAGTTAACT

>Seq84_4_64316_0.0001

GCTATGTGAGATTAACCCTTTAAAGTTAACT

>Seq85_4_64316_0.0001

GCTATGTGGAGATTCCCTTTAAAGTTAACT

>Seq86_4_64316_0.0001

GCTATGTGAGATTCTTTAAAGTTAACT

>Seq87_4_64316_0.0001

GCTAAGTGAGATTCCCTTTAAAGTTAACT

>Seq88_4_64316_0.0001

GCTATGTGAGATTCCCTAAAGTTAACT

>Seq89_4_64316_0.0001

GCAATGTGAGATTCCCTTTAAAGTTAACT

>Seq90_4_64316_0.0001

GCTAATGTGAGATTCCCTTTAAAGTTAACT

>Seq91_4_64316_0.0001

GCTTATGTGAGATTCCCTTTAAAGTTAACT

>Seq92_4_64316_0.0001

GCTATGTGAGATTGCCTTTAAAGTTAACT

>Seq93_3_64316_0.0000

GGTATGTGAGATTCCCTTTAAAGTTAACT

>Seq94_3_64316_0.0000

GCTATGTGGATTCCCTTTAAAGTTAACT

>Seq95_3_64316_0.0000

GCTATGTGAGATTCCCTTTGAAGTTAACT

>Seq96_3_64316_0.0000

GCTATGGTGAGATTCCCTTTAAAGTTAACT

>Seq97_3_64316_0.0000

GCTATGTGAGATCCCTTTAAAGTTAACT

>Seq98_3_64316_0.0000

GCTATGTGAAGATTCCCTTTAAAGTTAACT

>Seq99_3_64316_0.0000

GCTATGTGAGATTCCCTTTAAAGGTAACT

>Seq100_2_64316_0.0000

GCTATGTGAGATTCCCTTTAAAGTTATCT

>Seq101_2_64316_0.0000

GCTATGTTGAGATTCCCTTTAAAGTTAACT

>Seq102_2_64316_0.0000

GCTATGTGAGATTCCCTTTAAATTTAACT

>Seq103_2_64316_0.0000

GCTATGTGCGATTCCCTTTAAAGTTAACT

>Seq104_2_64316_0.0000

GCTATGTGAGATTCCCCTCAAAGTTAACT

>Seq105_2_64316_0.0000

GCTATGTGAGATTCCCTTTCAAGTTAACT

>Seq106_2_64316_0.0000

GATATGTGAGATTCCCTTTAAAGTTAACT

>Seq107_2_64316_0.0000

GCTATGTGAGATGCCCTTTAAAGTTAACT

>Seq108_2_64316_0.0000

GCTATGTGAGAGTCCCTTTAAAGTTAACT

>Seq109_2_64316_0.0000

GCTATGTGAGATTCCCTTTAAAGTTACCT

>Seq110_2_64316_0.0000

GCGATGTGAGATTCCCTTTAAAGTTAACT

>Seq111_2_64316_0.0000

GCTATGTGAGATTCCCCTTAAAAGTTAACT

>Seq112_2_64316_0.0000

GCTATGTGAGATTCCCTTGAAAGTTAACT

>Seq113_2_64316_0.0000

GCTATGTGAGATTAACCTTTAAAGTTAACT

>Seq114_2_64316_0.0000

GCTATGTGAGATTCCCTTTTAAAGTTAACA

>Seq115_2_64316_0.0000

TATGTGAGATTCCCTTTAAAGTTAACT

>Seq116_1_64316_0.0000

GCTATGTGAGATTCCAAAAAAGTTAACT

>Seq117_1_64316_0.0000

GCTATGTGAGATTGCCCTTTAAAGTTAACT

>Seq118_1_64316_0.0000

GCTATGTGAGATTCCCTGAAAGTTAACT

>Seq119_1_64316_0.0000

GCTATGTGAGATTCCCAAAGTTAACT

>Seq120_1_64316_0.0000

GCTATGTGAGATTCCCTTTAACGTTAACT

>Seq121_1_64316_0.0000

GCTATGAGATTCCCTTTAAAGTTAACT

>Seq122_1_64316_0.0000

GCTATGTGAGATTCTCTTCAAAGTTAACT

>Seq123_1_64316_0.0000

GCTATGTGAGATTCCCTCATAAAGTTAACT

>Seq124_1_64316_0.0000

CCTATGTGAGATTCCCTTTAAAGTTAACT

>Seq125_1_64316_0.0000

GCTATGTGCGATTCCCTTTAAAGTTCACT

>Seq126_1_64316_0.0000

GCTATGTGAGATTCCTTTAAAGTTAATT

>Seq127_1_64316_0.0000

GCTATGTGAGATTCCCTTTAAAGTTAAGT

>Seq128_1_64316_0.0000

GCTATGTGATTCCCTTTAAAGTTAACT

>Seq129_1_64316_0.0000

GCTATGTAAGATTCCTTTAAAGTTAACT

>Seq130_1_64316_0.0000

GCTATGTGAGATTCCCTTTAAAGTTAAACT

>Seq131_1_64316_0.0000

GCTATGTGAGATCTCCTTTAAAGTTAACT

>Seq132_1_64316_0.0000

GCTATGTGAGCTTCCCTTTAAAGTTAACT

>Seq133_1_64316_0.0000

CTTGTGAGATTCCCTTTAAAGTTAACT

>Seq134_1_64316_0.0000

GCTATGTGAGATTCCATTTAGAGTTAACT

>Seq135_1_64316_0.0000

GCTATGTGAGATTCCCTCCAAAGTTAACT

>Seq136_1_64316_0.0000

GCTCTGTGAGATTCCCTTTAAAGTTAACA

>Seq137_1_64316_0.0000

TGTGAGATTCCCTTTAAAGTTAACT

>Seq138_1_64316_0.0000

GCTAGGTGAGATTCCCTTTAAAGTTAACT

>Seq139_1_64316_0.0000

GATGTGAGATTCCCTTTAAAGTTAACT

>Seq140_1_64316_0.0000

GGCTGTGTGAGATTCCCTTTAAAGTTAACT

>Seq141_1_64316_0.0000

GCTATGTGAGATTAAAAAAAAGTTAACT

>Seq142_1_64316_0.0000

GCTATGTGAGATAACCCTTTAAAGTTAACT

>Seq143_1_64316_0.0000

CTATGTGAGATTCCCTTTAAAGTTAAC

>Seq144_1_64316_0.0000

GCTGTGAGATTCCCTTTAAAGTTAACT

>Seq145_1_64316_0.0000

GCTATGTGAGATTCCCTTTCAAGTTAACTC

>Seq146_1_64316_0.0000

GCTATGTGGGATTCCATTTAAAGTTAACT

>Seq147_1_64316_0.0000

GCTATGTGAGATTCCCTTTAAAGTTAACG

>Seq148_1_64316_0.0000

GCTATATGAGATTCCCTTTAAAGTTAACCT

>Seq149_1_64316_0.0000

GCTATGTGAGATTCCCTTTAAAGTGAACT

>Seq150_1_64316_0.0000

GCGTGAGATTCCCTTTAAAGTTAACT

>Seq151_1_64316_0.0000

GCTATGTGAGATTTTAAAGTTAACT

>Seq152_1_64316_0.0000

GCTATGTGAGATTCCCTTTAAAGTAACT

>Seq153_1_64316_0.0000

GCTATGTGAGATTCTCTTTAAAGTTAAAT

>Seq154_1_64316_0.0000

GCTATGTCAGATTCCCTTTAAAGTTAACT

>Seq155_1_64316_0.0000

GCTATGTGAGATTCCCCCTTAAAGTTAACT

>Seq156_1_64316_0.0000

GTTATGTGAGATTCCCCTTTAAAGTTAACT

>Seq157_1_64316_0.0000

GCTATGTGAGATTCCCTTTACAGTTAACT

>Seq158_1_64316_0.0000

GCTATGTGGGATTTCCCTTTAAAGTTAACT

>Seq159_1_64316_0.0000

GCTATGTGAGTTTCCTTTTAAAGTTAACT

>Seq160_1_64316_0.0000

GCTGTGTGTGATTCCCTTTAAAGTTAACT

>Seq161_1_64316_0.0000

GCTATGTGAGATTCCCTGTAAAGTTAACT

>Seq162_1_64316_0.0000

GCTATGGAGATTCCCTTTAAAGTTAACT

>Seq163_1_64316_0.0000

GCTATGTGAGATTTCCTTCAAAGTTAACT

>Seq164_1_64316_0.0000

GCTATGTGAGATTCCCTTTAAAAGTTAGCT

>Seq165_1_64316_0.0000

GCTATGTAGATTCCCTTTAAAGTTAACT

>Seq166_1_64316_0.0000

GCTATGTGAGATTCCCTTTAAAGTTAACTCCGCTCGAGCGGTGAATTGGTATCGACCATTTGTCGAGACTTTGCATCGTGCTATGTGAGATTCCCTTTAAAGTTAACT

>Seq167_1_64316_0.0000

CTATGTGAGATTCCCTTTTAAGTTAACT

>Seq168_1_64316_0.0000

GCTATGTGAGGATTTCCTTTAAAGTTAACT

>Seq169_1_64316_0.0000

GTTATGTGAGATTTCCTTTAAAGTTAACT

>Seq170_1_64316_0.0000

GCTATGTGAGATTCCGTTTAAAGTTAACT

>Seq171_1_64316_0.0000

GCTATGTGAGATTCCTTTTAAAGATAACT
